# Redistribution of centrosomal proteins by centromeres and Polo kinase controls partial nuclear envelope breakdown in fission yeast

**DOI:** 10.1101/2020.12.01.406553

**Authors:** Andrew J. Bestul, Zulin Yu, Jay R. Unruh, Sue L. Jaspersen

**Affiliations:** Stowers Institute for Medical Research, Kansas City, MO; Department of Molecular and Integrative Physiology, University of Kansas Medical Center, Kansas City, KS

**Keywords:** nuclear envelope breakdown, Polo kinase, LEM domain, Sad1-UNC-84 domain, spindle pole body

## Abstract

Proper mitotic progression in *Schizosaccharomyces pombe* requires partial nuclear envelope breakdown (NEBD) and insertion of the spindle pole body (SPB – yeast centrosome) to build the mitotic spindle. Linkage of the centromere to the SPB is vital to this process, but why that linkage is important is not well understood. Utilizing high-resolution structured illumination microscopy (SIM), we show that the conserved SUN-domain protein Sad1 and other SPB proteins redistribute during mitosis to form a ring complex around SPBs, which is a precursor for localized NEBD and spindle formation. Although the Polo kinase Plo1 is not necessary for Sad1 redistribution, it localizes to the SPB region connected to the centromere, and its activity is vital for redistribution of other SPB ring proteins and for complete NEBD at the SPB to allow for SPB insertion. Our results lead to a model in which centromere linkage to the SPB drives redistribution of Sad1 and Plo1 activation that in turn facilitate partial NEBD and spindle formation through building of a SPB ring structure.

## Introduction

Microtubule-organizing centers (MTOCs) are found throughout eukaryotes and carry out a vast array of cellular processes, including microtubule nucleation (Bettencourt-Dias and Glover, 2007; Wu and Akhmanova 2017; Kollman et al 2011). During mitosis, MTOCs known as the centrosome (metazoans) or spindle pole body (SPB, fungi) serve as the poles of the mitotic spindle as part of the spindle apparatus that facilitates accurate chromosome segregation. Failure of the centrosome/SPB to properly assemble the mitotic spindle results in chromosome missegregation and genomic instability (Nigg 2002; Ganem et al. 2009; Gönczy et al. 2015). In metazoans and in the fission yeast, *Schizosaccharomyces pombe*, the centrosome/SPB is located on the cytoplasmic surface of the nuclear envelope (NE) throughout interphase. As cells enter mitosis, nuclear envelope breakdown (NEBD) begins beneath the centrosome/SPB. In contrast to the complete NEBD of most metazoan cells, fission yeast restricts NEBD to the region localized only underneath the SPB (McCully and Robinow 1971, Tanaka and Kanbe 1986). This partial NE fenestration allows for SPB insertion into the nuclear membrane, which enables microtubules to access chromosomes to create the mitotic spindle (Ding et al 1997; Uzawa et al 2004; Cavanaugh and Jaspersen 2017). Partial NEBD is also observed in *Caenorhabditis elegans* early embryos, in the syncytial embryonic division cycles of *Drosophila melanogaster* and in numerous other mitotic systems, particularly in lower eukaryotes (Hachet et al. 2007; Portier et al. 2007; Paddy et al. 1996; Heath and Heath 1976).

The physical mechanism of NEBD has been most extensively examined in metazoans, which generally undergo a complete breakdown of the NE upon entry into mitosis. Critical regulators of NEBD in metazoans include cyclin dependent kinase 1 (CDK1/Cdc2), Polo-like kinase 1 (PLK-1) and Aurora B kinase. Beginning in prometaphase, these kinases act on targets such as nucleoporins, lamins and other membrane-associated proteins to bring about disassembly of nuclear pore complexes (NPCs), the nuclear lamina and the NE itself (reviewed in Lindqvist et al., 2009). Although fungi do not undergo complete NEBD and lack lamins, aspects of this mitotic NEBD and its regulation are conserved. For example, in the filamentous fungus *Aspergillus nidulans*, phosphorylation of nucleoporins regulates NPC remodeling needed for mitotic spindle formation within the intact NE (De Souza et al., 2004), while in fission yeast, mitotic upregulation of lipid synthesis is required for NE expansion (reviewed in Zach and Prevorovsky 2018).

A major unresolved question is how NE remodeling is spatially regulated in cells that undergo partial NEBD. This question has been extensively studied in fission yeast meiosis where the linkage of a cluster of telomeres (called the telomere bouquet) to SPB through the NE is vital to trigger partial NEBD at the SPB, which is needed for SPB insertion into the NE and spindle formation (Tomita and Cooper, 2007; Klutstein and Cooper, 2014). The SPB receptor for telomeres during meiosis is the LINC (Linkage of Nucleoskeleton and Cytoskeleton) complex, composed of the outer nuclear membrane (ONM) KASH-domain protein Kms1 and the inner nuclear membrane (INM) SUN-domain protein, Sad1. Telomeric linkage to the LINC is achieved by association of the meiosis-specific telomere-binding proteins Bqt1 and Bqt2 with Sad1 (Chikashige et al., 2006). Spatialized NEBD at the SPB is also regulated by chromosome-NE association in mitotic cells. Here, the meiosis-specific INM KASH-protein Kms1 is replaced by its mitotic ortholog Kms2 (Wälde and King, 2014; Bestul et al., 2017). Also, the telomere bouquet is replaced by centromeres, which are clustered underneath the SPB during interphase of mitotic cells through association of the mitotic centromere binding protein Csi1 with Sad1 (Funabiki et al., 1993; Hou et al., 2012). However, loss of Csi1 only partially disrupts Sad1-centromere linkage, implicating other unknown linkage factors such as the LEM-domain proteins Lem2 and/or Man1 that also bind to centromeres/telomeres (Hiraoka et al., 2011; Steglich et al., 2012). A single centromere bound to the LINC complex is sufficient for NE remodeling, SPB insertion and spindle formation; in *sad1.2*, which displays a partial loss of centromere-binding function, cells in which all three centromeres detach from the SPB have a defect in mitotic progression (Fennell et al., 2015; Fernandez-Alvarez et al., 2016). Analysis of *sad1.2* cells showed a defect in NEBD, providing evidence that chromosome linkage via the LINC complex regulates partial NEBD at mitotic entry (Fernandez-Alvarez et al., 2016). How a chromosome-NE linkage leads to partial NEBD and SPB insertion is unknown, although it is hypothesized that chromatin may increase the critical concentration of mitotic regulators at the SPB to bring about NE remodeling (Fernandez-Alvarez and Cooper 2017a). A leading candidate is cyclin-dependent kinase 1 (CDK1/Cdc2), which associates with the nuclear side of the NE underneath the SPB (Alfa et al., 1990).

The budding yeast SPB is inserted into the NE by the spindle pole insertion network (SPIN), which includes the SUN protein Mps3 and its binding partner, Mps2, that together form a non-canonical Linkage of Nucleoskeleton and Cytoskeleton (LINC) complex (Rüthnick et al., 2017; Chen et al., 2019). The SPIN forms a donut-like structure that anchors the SPB in the NE. The SPIN also may play a role in partial NEBD, possibly through localized changes in lipid composition, recruitment of NE remodeling factors such as Brr6 and/or alterations in NPCs (Friederichs et al., 2011; Rüthnick et al., 2017; Zhang et al., 2018). Like Mps3, Sad1 also localizes to a ring-like structure around duplicated SPBs at mitotic entry (Bestul et al., 2017), raising the possibility that Sad1 plays a key role in partial NEBD, SPB insertion and spindle assembly in fission yeast. Here, we investigate the link between Sad1, ring formation and centromere linkage using high-resolution structured illumination microscopy (SIM). We find that Sad1, Kms2, the mitotic regulator Cut12 and Cut11, a dual component of SPBs and NPCs all form a ring-like structure at the SPB that is Sad1- and centromere-dependent. This novel mitotic ring structure is required for partial NEBD mediated by Sad1-centromere binding, which recruits Polo kinase (plo1+) to the NE. Polo kinase, in turn, is needed to drive formation of the mitotic SPB ring that allows for partial NEBD and SPB insertion.

## Results

### Identifying components of the fission yeast mitotic SPB ring

Previously, we examined Sad1 localization using SIM in cells arrested at G2/M using the *cdc25.22* mutant (Bestul et al., 2017). We found images in which Sad1-GFP shifted from underneath the duplicated side-by-side SPBs to a full or partial ring surrounding the SPB core. This ring-like pattern of localization is reminiscent of Mps3 in budding yeast, which like other SPIN components, surrounds the SPB core (Chen et al., 2019).

To identify other SPB proteins that redistribute into a ring like Sad1, we introduced sixteen previously GFP-tagged SPB components (West et al., 1998; Bestul et al., 2017) into a *cdc25.22* strain containing Ppc89-mCherry to mark the SPB core. Cells were arrested at G2/M by growth at 36°C for 3.5 h then were released into mitosis by shifting to 25°C. Examination of cells at 0, 10, 20 and 30 min allowed us to follow ring formation and SPB insertion upon mitotic entry. Critically, Ppc89-mCherry and other components of the SPB core (GFP-Pcp1, Sid4-GFP) appeared as two foci at all time points, indicating that the SPB core does not reorganize upon entry into mitosis (Fig. S1A). Consistent with observations that core proteins do not form a ring, electron microscopy (EM) shows that the laminar SPB core is simply lowered into a fenestrated region of the NE during mitosis (McCully and Robinow 1971, Tanaka and Kanbe 1986; Ding et al., 1997; Uzawa et al., 2004).

Four proteins redistributed into ring-like structures: Sad1, Kms2, Cut12 and Cut11 (Fig 1A). Sad1 and Cut11 rings are robust, encompassing almost the entire perimeter of duplicated SPBs, while Cut12 and Kms2 only form partial rings. Cut11 and Kms2 localization to the region surrounding the SPB is not unexpected as both contain transmembrane domains and are orthologous (Cut11) or similar (Kms2) to the SPIN components Ndc1 and Mps2 that distribute around the SPB in budding yeast (Rüthnick et al., 2017; Chen et al., 2019). However, Cut12 does not have a budding yeast ortholog and it lacks a transmembrane domain. We hypothesize that Cut12 is targeted to a ring through its interaction with Kms2 in much the same way that the soluble protein Bbp1 is targeted to the SPIN by Mps2 (Wälde and King, 2014; Kupke et al., 2017; Schramm et al., 2000). Kms2-dependent Cut12 targeting could explain why these rings are similar in appearance but different than fuller rings formed by Sad1 and Cut11.

**Figure 1.**
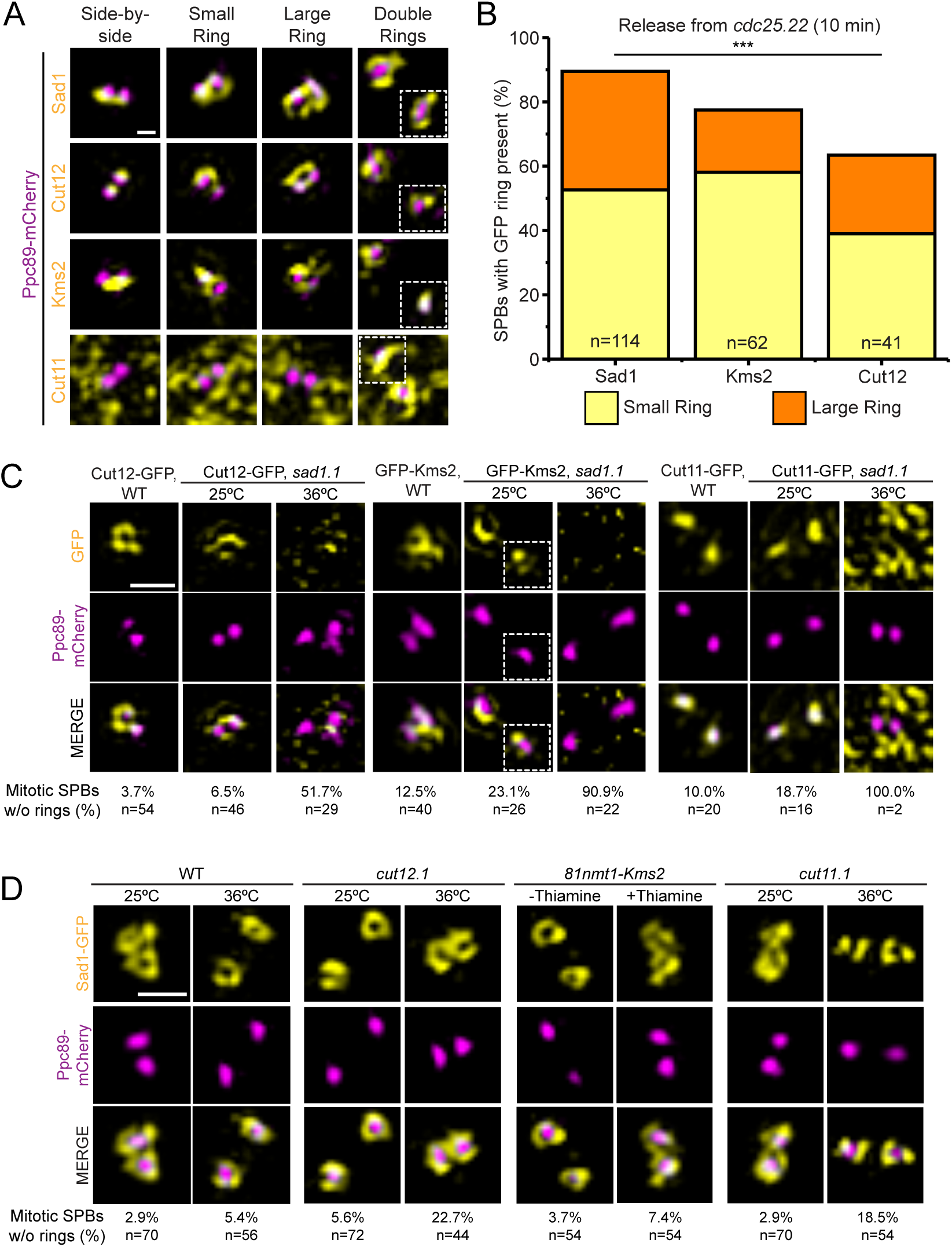
Four SPB proteins constitute the Sad1-dependent *S. pombe* mitotic SPB ring. (A-B) *cdc25.22* cells containing Ppc89-mCherry and the indicated GFP-tagged SPB components were arrested at G2/M by growth at 36°C for 3.5 h, then released into mitosis by shifting back to 25°C. Samples were taken at 0, 10, 20 and 30 min for analysis by SIM. (A) Example images of Sad1-GFP, Cut12-GFP, GFP-Kms2 and Cut11-GFP (yellow) relative to the SPB (magenta). At the arrest, SPB rings have not formed (side-by-side); next, a small ring is observed; followed by a large ring that encapsulates both SPBs; finally, two separated (double) rings are observed as the poles move apart (one SPB in the same cell is shown as inset, white boxes). The punctate background of Cut11-GFP is due to its localization to NPCs, which occurs throughout the cell cycle (West et al., 1998). Bar, 200 nm. (B) Percentage of SPBs containing small (yellow) and large (orange) rings of the indicated protein was quantitated at 10 min after *cdc25.22* release. Cut11-GFP does not localize to the SPB at this time point so it was not included. ***p=0.0013 based on χ^2^ test. (C) SIM images from wild-type or *sad1.1* cells containing Ppc89-mCherry (magenta) and the indicated GFP-tagged SPB component (yellow) grown at 25°C or shifted to 36°C for 3.5 h. Bar, 500 nm. Percentage of SPBs with no GFP signal in a ring formation is indicated below each corresponding column along with the number of SPBs analyzed (n). (D) SIM images from *cut11.1* and *cut12.1* strains containing Sad1-GFP (yellow) and Ppc89- mCherry (magenta) grown at 25°C or shifted to 36°C for 3.5 h, and from *81nmt-HA- Kms2* cells with Sad1-GFP (yellow) and Ppc89-mCherry (magenta) grown in the absence or presence of 20 μM thiamine for 16 h, which was previously shown to deplete *HA-Kms2* (Walde and King, 2014). Bar, 500 nm. Percentage of SPBs no Sad1- GFP ring formation is indicated below each corresponding column along with the number of SPBs analyzed (n).

To ensure that ring formation is not an artifact caused by the prolonged G2/M arrest in *cdc25.22*, we examined asynchronously growing cells containing GFP-tagged versions of the four identified ring proteins and Ppc89-mCherry (Fig. S1A). All four proteins rearrange to surround the SPB core at mitotic entry. Thus, ring formation is a natural phenomenon and is not an artifact caused by cell cycle manipulation using the *cdc25.22* allele. The rings formed in asynchronous cells are often smaller and more difficult to identify than those found in *cdc25.22* cells, so we utilized the G2/M arrest throughout this study to maximize ring detection and to provide information about the kinetics of ring assembly. Ring formation can be divided into four distinct steps: 1) in G2/M, SPBs are in a side-by-side configuration with protein present at or between the SPBs; 2) as cells enter into mitosis, a small ring is observed that has a diameter similar in size to a single SPB; 3) later, the small ring expands into a large ring that encompasses both SPBs; 4) finally, as SPBs separate, a ring surrounds each of the two SPBs (double rings) (Fig 1A).

### Stepwise formation of the SPB ring is initiated by Sad1

Quantitation of the fraction of cells containing at least one observable ring at 10 min after *cdc25.22* release pointed to a temporal hierarchy of ring formation beginning with Sad1. At 10 min, 89.5% of Sad1 had redistributed to a ring, compared to 77.5% of Kms2 and 63.4% of Cut12 (Fig 1B). In addition, 36.9% of the Sad1 rings were in the more mature large ring conformation at this time, compared to 19.2% of Kms2 and 24.4% of Cut12. This suggests that Sad1 reorganization precedes that of Cut12 and Kms2. Examination of *cdc25.22* cells containing Sad1-mCherry and Cut12-GFP confirmed this idea; 10 min after release, 72.2% of cells had both a Cut12 and Sad1 ring, 27.8% of the cells had only a Sad1 ring present and no cells had only a Cut12 ring (Fig S1B). Cut11-GFP, which is found at NPCs throughout the cell cycle (West et al., 1998), localized to double rings at or near the time of SPB separation, approximately 30 min following release from *cdc25.22* or during prometaphase (Fig 1A). In a few rare cases, we observed Cut11-GFP large ring structures prior to SPB separation. As Cut11- GFP small rings were never seen, it seems that Cut11 ring formation is not regulated at mitotic entry like the other three proteins observed. From this data, we propose an order of ring formation that starts with Sad1, followed by Kms2 and Cut12 and then finally Cut11 right before SPB separation.

Based on our temporal model of ring assembly, Sad1 serves as the gatekeeper. Loss of *sad1+* function should lead to defects in redistribution of other components. Indeed, in *sad1.1* mutants at 36°C, 51.7% to 100% of cells did not have Cut12-GFP, GFP-Kms2 or Cut11-GFP rings at mitotic SPBs compared to 3.7% to 12.5% ring loss seen in wild-type cells grown at the same temperature (Fig 1C). This data suggests that Sad1 regulates ring distribution of Kms2 and Cut12 at the SPB. Because Cut11 only localizes to rings in late prometaphase or early metaphase (Fig 1A), *sad1.1* cells do not progress sufficiently into mitosis for Cut11 to localize to the SPB and form rings (West et al., 1998). Therefore, we cannot determine if Cut11 ring formation is Sad1-dependent or Sad1- independent.

As the gatekeeper, we also predict that Sad1 redistribution to a ring would be unaffected by loss of function in the downstream components. At 36°C, 5.4% of mitotic SPBs in wild-type cells did not have a Sad1-GFP ring compared to 22.7% of *cut12.1* mutants, 7.4% of *81nmt1-HA-Kms2* mutants and 18.5% of *cut11.1* mutants (Fig 1D). The fact that *cut12.1* mutants have a higher fraction of cells without a Sad1-GFP ring or that many Sad1-GFP rings, particularly in *cut11.1* mutants, were not as full or complete as in wild-type cells may indicate that Cut12 and/or Cut11 stabilizes the Sad1-GFP ring. This is consistent with the preferential association of Sad1 with one SPB, as previously described using widefield microscopy (Bridge et al., 1998; Tallada et al., 2009). However, in our experiments, Sad1-GFP rings are typically visible at both SPBs in all the mutants, consistent with the idea that Sad1 ring formation is independent of Kms2, Cut12 and Cut11 function.

### Formation of the mitotic ring coincides with partial NEBD

The appearance of Sad1, Kms2, Cut12 and Cut11 in ring-like structures in prometaphase coincides with the timing of SPB insertion, which requires creation of the NE pore known as a fenestra. In wild-type cells, a small part of the NE is broken down and is quickly plugged by the SPB, keeping the NE intact during mitosis (Ding et al., 1997; Uzawa et al., 2004). An NLS-GFP reporter (*nmt1-3x-NLS-GFP*) to monitor nuclear integrity remains in the nucleus at all times in wild-type cells (Tallada et al., 2009; Fernandez-Alvarez et al., 2016; Tamm et al., 2011). If SPB insertion is disrupted (*brr6.ts8*), large holes in the NE form (Tamm et al., 2011), and the ratio of nuclear to cytoplasmic (N:C) GFP of the reporter decreases (Fig 2A).

**Figure 2.**
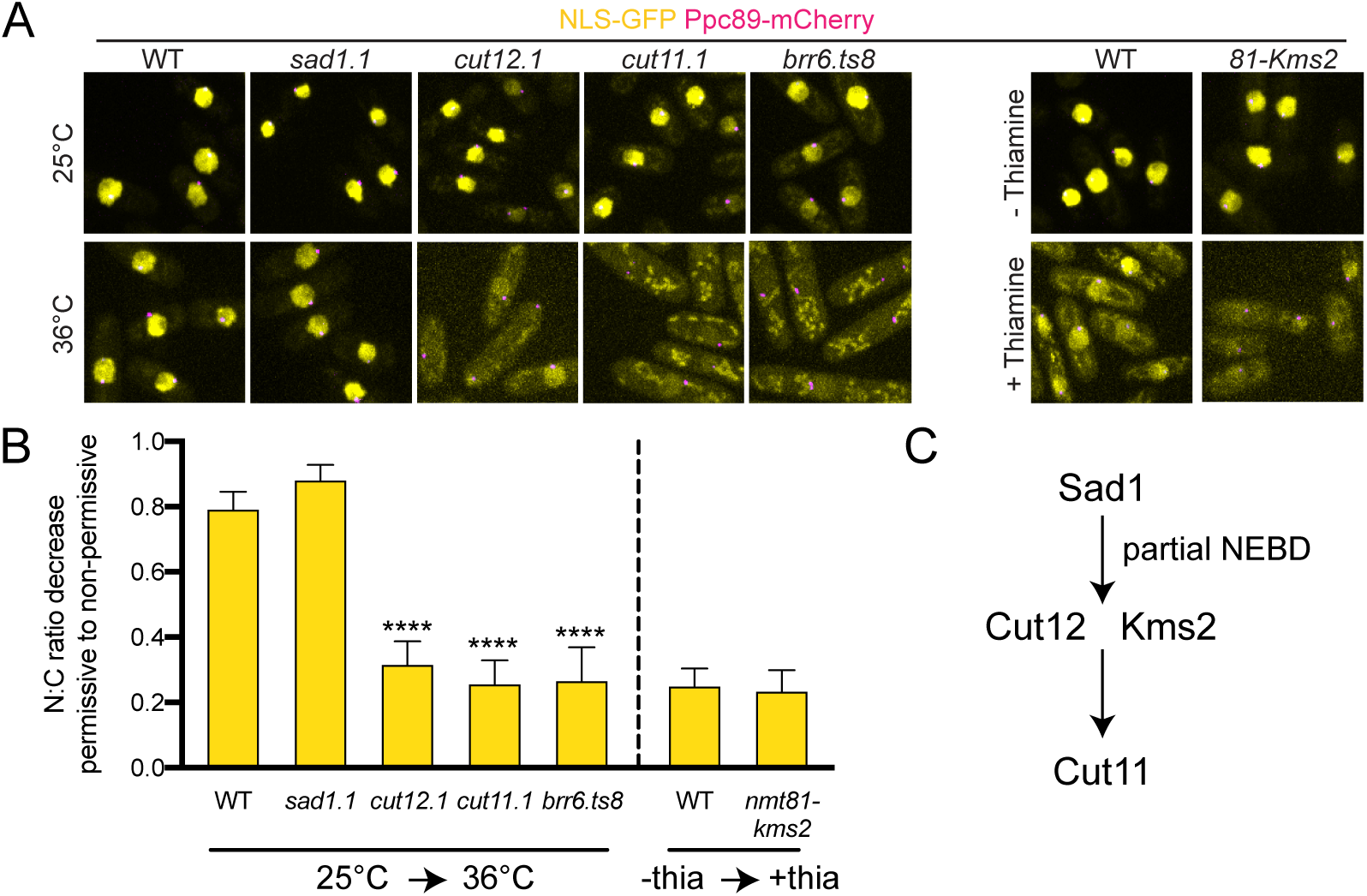
Nuclear integrity is maintained in the absence of *sad1+* function. The soluble nuclear reporter nmt1-3x-NLS-GFP (yellow) was introduced into wild-type and mutant strains containing Ppc89-mCherry (magenta) to test NE integrity. Cells were grown at 25°C or shifted to 36°C for 3.5 h (for wild-type; *sad1.1, cut12.1, cut11.1* and *brr6.ts8*) or grown for ∼16 h in the absence or presence of 20 μM thiamine (for WT and *81nmt-HA-Kms2*). (A) Representative confocal images. Bar, 5 μm. (B) The ratio of total nuclear (N) to cytoplasmic (C) GFP fluorescence was determined for each strain and a normalized ratio based on the permissive condition was determined. Plotted is the decrease of each N:C ratio when going from the permissive condition (25°C or no thiamine) to the restrictive condition (36°C or 20 μM thiamine). A value of 1.0 would indicate no decrease between the two conditions. Errors bars, SEM. ****p<0.001, based on χ^2^ test compared to wild-type. n=100. (C) Partial NEBD timing relative to ring formation based on data shown here, which is consistent with mutant data previously reported (Tallada et al., 2009; Fernandez-Alvarez et al., 2016; Tamm et al., 2011).

If the mitotic ring is involved in partial NEBD during SPB insertion, it seemed likely that blocking ring assembly by mutation of *sad1+*, *kms2+*, *cut12+* or *cut11+* might uncouple fenestration and SPB insertion, leading to alterations in NE integrity. Previous reports indicate this is indeed true (Tallada et al., 2009; Fernandez-Alvarez et al., 2016; Tamm et al., 2011), however, it is was important to directly analyze the same NLS-GFP reporter in *sad1.1, cut12.1, cut11.1* and *81nmt1-HA-Kms2* mutants so we could directly compare effects on NE integrity. From this side-by-side comparison, we observed a range of phenotypes that can be categorized as: 1) severe/complete loss of NE integrity, 2) partial loss of NE integrity and 3) no loss of NE integrity (Fig 2B). Similar to the *brr6.ts8* (N:C 1.4±0.1) mutant, *cut11.1* (N:C 1.5±0.1) mutants showed complete loss of NE integrity. *cut12.1* and *81nmt-HA-Kms2* cells showed a partial loss of NE integrity with N:C ratios of 2.2±0.1 and 2.3±0.1 under non-permissive conditions, values that were significant compared to wild-type controls.

In contrast, even at restrictive conditions, the N:C ratio for *sad1.1* mutants remained high (9.5±0.3), similar to wild-type cells (10.2±0.3). Fernandez-Alvarez et al., 2016 also found that the NE remained intact in cells carrying the *sad1.2* allele (Fig 2A-B). Thus, *sad1+* function is required for NEBD. Without Sad1, other components do not localize (Fig 1C) and further steps leading to NEBD cannot occur. Cut12, Kms2 and Cut11 are not required to initiate NEBD as partial or total loss of NE integrity occurs in their absence (Fig 2C). Rather, they play roles in SPB insertion downstream. Our model that Sad1 is the first protein to redistribute into a ring and that its relocalization is needed for Cut12 and Kms2 ring formation and partial NEBD is consistent with cytological and genetic data showing that Sad1 interacts with both Kms2 and Cut12 (Walde and King 2014; Miki et al., 2004). Although Sad1 also interacts with Cut11 in the yeast two hybrid system (Varberg et al., 2020), no physical interactions between Cut11 and ring components have been reported in fission yeast, supporting the possibility that Cut11 recruitment may occur later in prometaphase in a Sad1-independent pathway.

### Centromeric-SPB linkage proteins Lem2 and Csi1 contribute to Sad1 redistribution into the mitotic SPB ring

As the gatekeeper whose function is needed to trigger partial NEBD, we were interested in determining how Sad1 is regulated. Sad1 redistribution typically begins by the formation of a small single ring, which we assumed surrounded either the ‘old’ SPB present from the previous cell cycle or the ‘new’ SPB formed by SPB duplication. Detailed inspection of the Sad1-GFP single ring showed it often formed between the two SPBs under the bridge region (18/38; non-SPB) rather than surrounding one of two SPBs (20/38; SPB) (Fig S2A). This suggests that the Sad1 redistribution is not linked to the SPB per se but to extrinsic landmarks, such as the centromere that is attached to the SPB through a Sad1-based LINC complex (Fernandez-Alvarez et al., 2016). Consistent with this idea, we observed that the Sad1 rings are positioned near the centromere, which was marked with Mis6-GFP (Fig S2B).

The coincidence of Sad1 rings near the centromere suggested that centromeric proteins might regulate Sad1 redistribution. Two candidates stood out as possible regulators: Csi1, a Sad1-interacting protein whose loss leads to partial disruption of the centromere-SPB linkage (Hou et al., 2012); and Lem2, which localizes to the SPB throughout interphase and early mitosis (Hiraoka et al., 2011), binds to chromatin near the centromere and functions in NE reformation (Gu et al., 2017; Barrales et al., 2016; Banday et al., 2016). A double deletion of *lem2+* and *csi1+* (*lem2Δ csi1Δ*) leads to a defect in mitotic spindle formation presumably due to loss of centromere-SPB tethering (Fernandez-Alvarez and Cooper 2017b).

To determine if Csi1 and/or Lem2 regulates Sad1, we examined the distribution of Sad1-GFP in *csi1Δ*, *lem2Δ* and *lem2Δ csi1Δ* backgrounds in normal (25°C) and stressed (36°C) conditions, which previously led to growth and nuclear morphology defects in a number of deletion mutants in INM components (Hiraoka et al., 2011). As a control, we also examined Sad1-GFP in other deletion strains for proteins involved in centromere or NE-binding and/or NE reformation: Nur1 (Banday et al., 2016); Bqt4 (Hirano et al., 2018; Hu C et al., 2019); Cmp7 (Gu et al., 2017) and Ima1 (Hiraoka et al., 2011; Steglich et al., 2012). The only single gene deletion mutant to significantly affect Sad1 ring formation was *csi1Δ*, where we observed a 29.6% reduction in ring formation at 36°C (Fig 3A-B, Fig S2C-D). The nature of the temperature dependence is unknown; it could be linked to temperature-dependent changes in lipid composition, protein folding, stress response pathways or other factors. Although *lem2Δ* did not have a phenotype on its own (Fig S2C-D), its loss further exacerbated *csi1Δ* resulting in a 36.1% decrease in Sad1-GFP ring formation in *lem2Δ csi1Δ* mutants at 36°C as well as a defect at 25°C (Fig 3A-B). This suggests that Csi1 and to a lesser degree Lem2 play a role in Sad1 ring formation.

**Figure 3.**
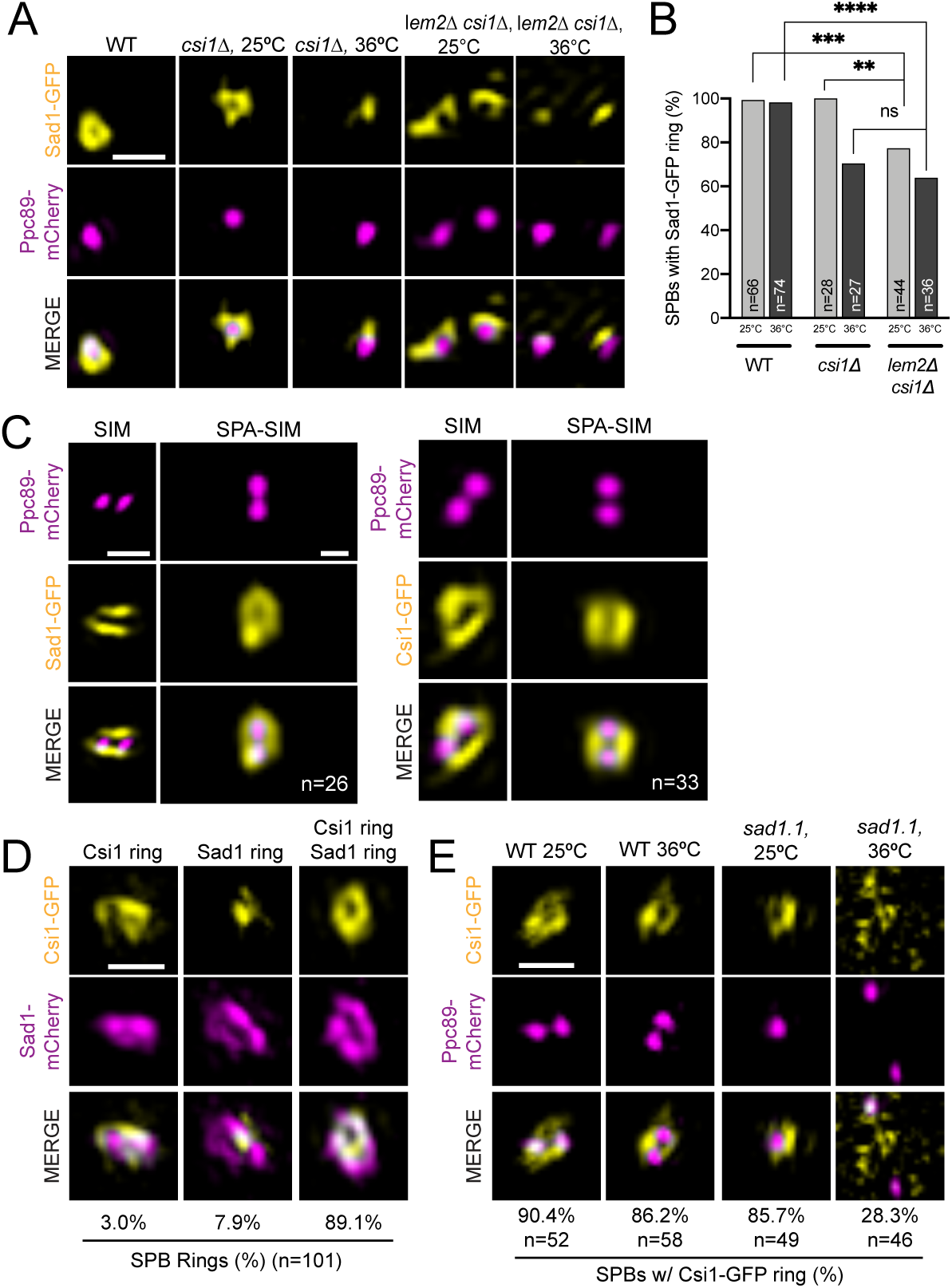
Centromeric protein Csi1 forms a mitotic ring and regulates Sad1 mitotic SPB ring formation. (A-B) Wild-type, *csi1Δ* or *lem2Δ csi1Δ* cells with Sad1-GFP (yellow) and Ppc89-mCherry (magenta) were grown for 4 h at 25°C or 36°C and then imaged. (A) Representative SIM images. (B) Percentage of SPBs with a Sad1 ring. P values were determined using the χ^2^ test, ns=not significant; **p=0.028; ***p=0.0056; ****p<0.001. Bars, 500 nm. (C) Individual SIM images of *cdc25.22* arrested cells containing Sad1- or Csi1-GFP (yellow) and Ppc89-mCherry (magenta) (left columns). Single particle averaging (SPA) was used to combine the indicated (n) number of individual SIM images (right columns). Bar, 500 nm. (D) SIM images of Sad1- mCherry (magenta) and Csi1-GFP (yellow) in *cdc25.22* arrested cells that were then released at 25°C for 10 min before imaging. Three configurations were observed in the indicated fraction of cells: Csi1 ring with no Sad1 ring (left), Sad1 ring with no Csi1 ring (center), and rings of both Sad1 and Csi1 (right). n=101. Bar, 500 nm. (E) SIM images of Csi1-GFP (yellow) and Ppc89-mCherry (magenta) in wild-type and *sad1.1* backgrounds, grown as described in Fig 1D. The percentage of cells containing a ring of Csi1-GFP around the SPB was determined for each based on the indicated number of cells (n).

Consistent with a role in Sad1 ring formation, we observed that both Csi1-GFP and Lem2-GFP formed ring-like structures similar to Sad1-GFP at high resolution (Fig 3C, S3A). Single particle averaging (SPA) using Ppc89-mCherry as a fiducial marker allowed us to align duplicated, but unseparated SPB pairs in G2/M cells over multiple SIM images (Burns et al., 2015; Bestul et al. 2017). From SPA-SIM analysis, we could compare the average protein distribution around the SPB for Sad1, Csi1 and Lem2 (Fig 3C, S3A). Csi1-GFP had a ring diameter slightly smaller than Sad1-GFP, whereas the diameter of the Lem2-GFP ring was larger, which was confirmed by measurements of ring diameter in individual cells (Fig S3B). Importantly, analysis of Csi1-GFP and Sad1-mCherry in synchronized cells (using *cdc25.22*) showed that Csi1 and Sad1 re- distribute into rings with similar timing (Fig 3D). Furthermore, *sad1.1* loss-of-function specifically blocked Csi1 ring formation (Fig 3E). In contrast, Lem2-GFP localizes to the SPB in interphase cells (Hiraoka et al., 2011), a time in the cell cycle when Sad1 does not form a ring (Bestul et al. 2017). Lem2-GFP persists in a ring at the SPB throughout prometaphase until SPB separation (signified by Cut11-GFP ring formation) (Fig S3C). During this stage of the cell cycle Lem2 and Sad1 rings co-exist, which may explain why Lem2-GFP ring formation was also impaired in a *sad1.1* background (Fig S3D) —Sad1 could be required to stabilize the Lem2 ring structure during prometaphase.

Alternatively, Lem2 may have begun its disassembly at the *sad1.1* arrest point. The size, timing and Sad1-dependence of Csi1 localization are consistent with a direct role for Csi1 in Sad1 ring formation and mitotic progression. While Lem2 may also be involved, our data suggest a more indirect role.

#### Attachment to the centromere is essential to Sad1 SPB ring formation

Previous work showed that *sad1.2* mutants exhibit a partial defect in centromere-binding that increases in severity with prolonged incubation at the non-permissive temperature of 36°C (Fernandez-Alvarez et al., 2016). To test if centromeres are directly involved in Sad1 reorganization at mitotic onset, we examined the ability of Sad1.2-GFP to form SPB rings. We hypothesized that ring formation would correlate with centromere-binding, which we tested by examining organization 4 h (partial loss) and 8 h (total loss) following shift to 36°C. At 25°C, 95.5% of cells contains a Sad1.2- GFP ring. This decreases to 78.9% if cells are shifted to 36°C for 4 h and further decreases to 36.5% after 8 h at 36°C (Fig 4A-B), strongly suggesting that centromere-binding is correlated with Sad1 ring formation. The observation that *csi1Δ* did not exacerbate Sad1.2-GFP ring formation at 36°C indicates that centromeric attachment is largely abolished due to the mutation in the *sad1.2* gene (Fig 4A-B). The finding that the *csi1Δ sad1.2* double mutant showed reduced ring formation at 25°C suggests that while Csi1 is a major linker for Sad1 and the centromere, it is not the only attachment factor. This is consistent with our and others’ data that *csi1Δ* is only partially penetrant (Hou et al., 2012; Fernandez-Alvarez and Cooper, 2017b).

**Figure 4.**
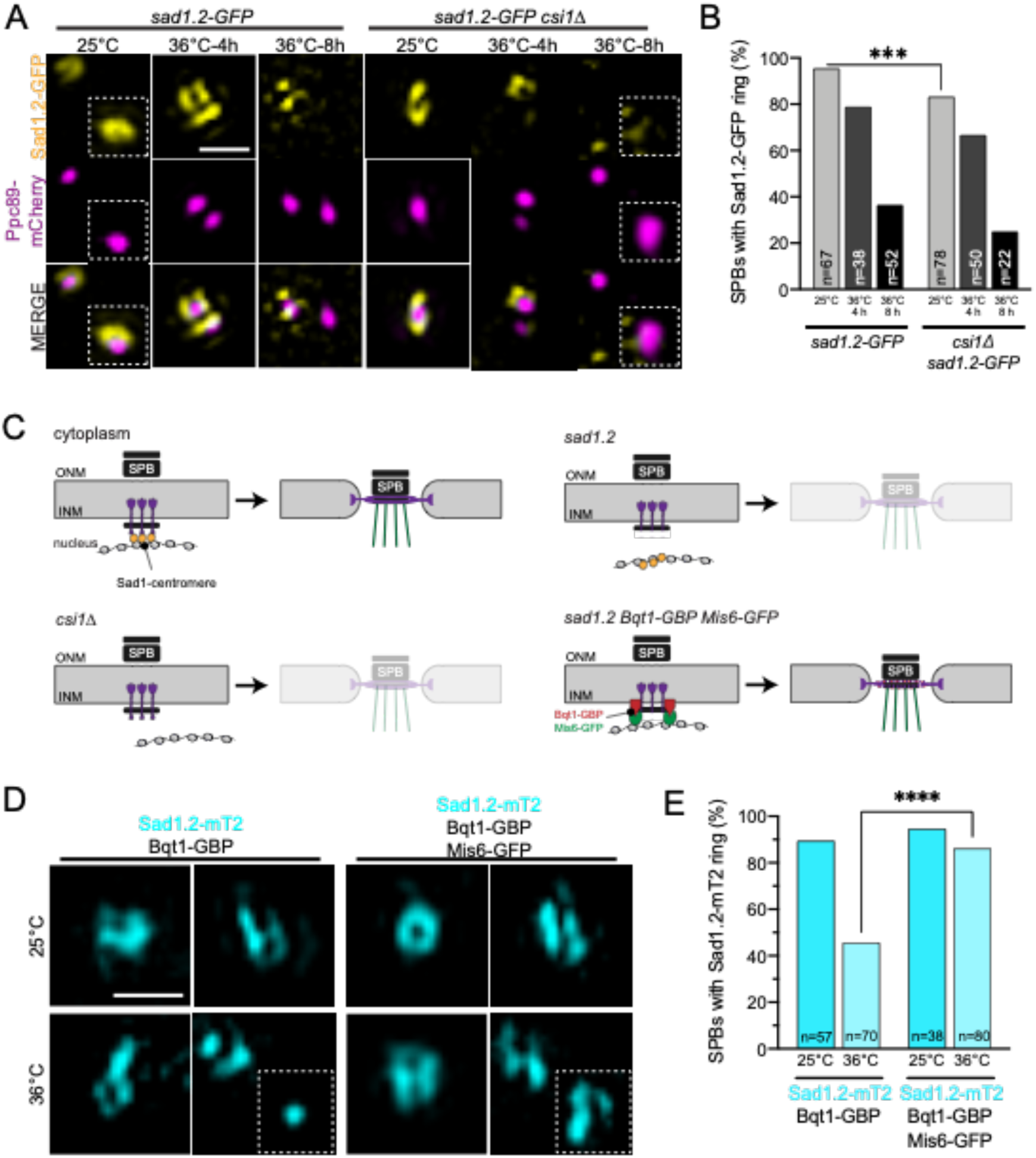
Sad1 attachment to the centromere is vital for Sad1 mitotic ring formation. (A-B) Sad1.2-GFP (yellow) in *csi1+* or *csi1Δ* cells with Ppc89-mCherry (magenta) were grown at 25°C for 4 h or at 36°C for 4 and 8 h before imaging. (A) Representative SIM images of Sad1.2-GFP rings at the SPB. The second SPB is shown in the inset. (B) Percentage of mitotic SPBs with a Sad1.2-GFP ring. P values were determined using the χ^2^ test. All are not significant, except ***p=0.007. (C) Schematic of centromere binding to the SPB. In wild-type cells (upper left), Sad1 (purple) is tethered by Csi1 (yellow) to the centromere, which leads to Sad1 ring formation at mitotic entry. Defects in the centromere attachment through *csi1Δ* (lower left) or *sad1.2* (upper right) disrupt Sad1 ring formation. To test if centromere attachment is sufficient to form the Sad1 ring, an artificial tethering system using Bqt1-GBP (lower right), which binds to both Sad1.2 and Mis6-GFP was used. (D-E) Sad1.2-mT2 Bqt1-GBP with and without Mis6-GFP were grown at 25°C for 4 h or at 36°C for 8 h. (D) Representative SIM images of Sad1.2-mT2 (cyan) mitotic rings. (E) Percentage of cells with Sad1.2- mT2 rings. P values were determined using the χ^2^ test. All are not significant, except ****p<0.001. Bars, 500 nm.

To confirm that centromere attachment alone was required for Sad1 reorganization, we rescued the Sad1.2 SPB ring formation defect by forcing centromere-SPB attachment using a previously described GFP-binding protein (GBP) fused to Bqt1, a meiotic protein that is still able to bind to the N-terminus of Sad1.2 (Fernandez-Alvarez et al., 2016). By adding GFP to a centromeric protein, Mis6 (Mis6-GFP), we can trigger centromere tethering: Mis6-GFP binds Bqt1-GBP, which then binds to Sad1.2 (Fig 4C). In the Bqt1- GBP background, Sad1.2-mTurquoise2 (mT2) SPB ring formation drops from 89.5% at 25°C to 45.7% at 36°C for 8 h, similar to Sad1.2-GFP levels seen above. However, when we force centromere attachment with Mis6-GFP in the Bqt1-GBP background, then Sad1.2-mT2 ring formation is fully rescued (Fig 4D-E). Collectively, these experiments show that centromere attachment alone is sufficient to drive Sad1 ring formation in mitotic cells. A key question is how centromeric tethering drives Sad1 reorganization. Using Sad1 distribution, we assayed factors involved in centromere-based signaling coincident with mitotic entry to test various potential regulators.

### Complete SPB ring formation and NEBD is regulated by Polo Kinase

Entry into mitosis is exquisitely regulated in most organisms to ensure that NEBD, spindle formation and chromosome segregation only occur once DNA replication has been completed. In fission yeast, a network of kinases and phosphatases control the G2/M transition (reviewed in Hagan IM, 2008), including the highly conserved CDK1, *cdc2+* (Nurse and Thuriaux, 1980); the Polo kinase, *plo1+* (Ohkura, Hagan and Glover, 1995); and Aurora B kinase, *ark1+* (Petersen et al., 2001; Leverson et al., 2002). In metazoans, different steps in NEBD are controlled by CDK1 and Polo kinase including phosphorylation of SUN1 (Patel et al., 2014). The role of kinases in Sad1 distribution in fission yeast is unknown, although the association of Cdk1 with centromeres during interphase has been proposed to underly the Sad1-centromere mediated NEBD and spindle formation (Fernandez-Alvarez and Cooper 2017a; Decottignies et al., 2001).

To inactivate *ark1+* and *plo1+*, we utilized temperature-sensitive strains, *ark1-T7* (Bohnert et al., 2009) and *plo1-24c* (Bähler et al., 1998), while for *cdc2+*, we utilized an analog-sensitive strain, *cdc2-asM17* (Aoi et al., 2014). Each of these kinase mutants were put into restrictive conditions for 4 h and then assayed for Sad1 ring formation (Fig 5A). *cdc2-asM17* had no effect on Sad1 ring formation as 96.4% of cells at the non-permissive condition formed Sad1 rings, which is very similar to wild-type levels (97.6%). To verify that *cdc2-asM17* was inactivated, we scored the percentage of mitotic cells in the population, based on Sad1 ring formation. In controls, ∼20% of cells were judged to be in mitosis; when *cdc2-asM17* was inhibited, the percentage of mitotic cells jumped to 77.5% (Fig S4). This indicates that the *cdc2-asM17* strain is blocked in mitosis at a step after Sad1 ring formation. *ark1-T7* mutants displayed a mild (83.8%) decrease in Sad1 ring formation at 36°C. Therefore, it is unlikely that Cdk1 or Ark1 is required for Sad1 ring formation. The most significant loss was seen with *plo1-24c*, which had only 38.5% Sad1 ring formation at 36°C (Fig 5A-B). This suggests that Polo kinase is required at a step of Sad1 reorganization.

**Figure 5.**
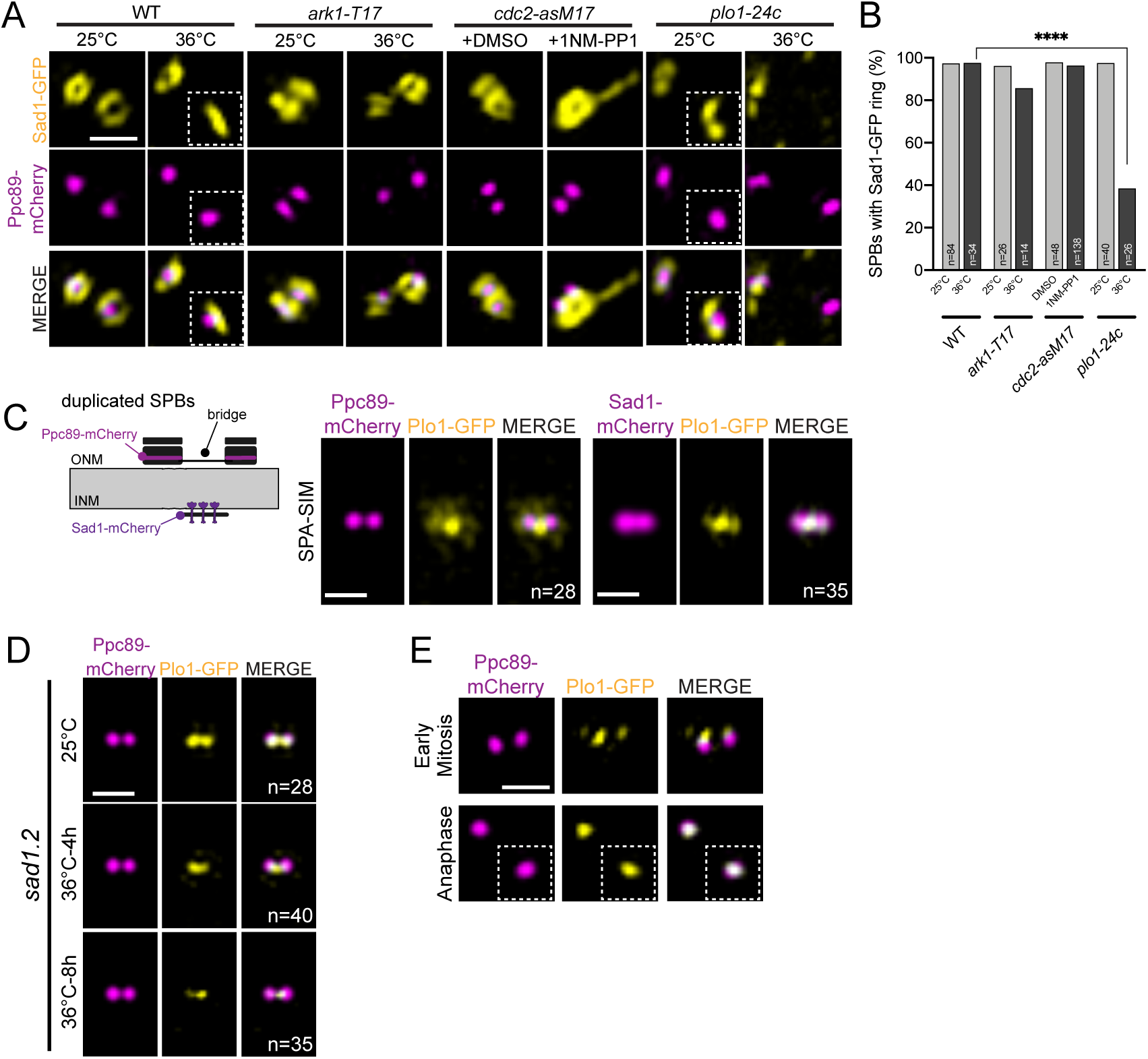
Polo kinase is necessary for Sad1 ring formation but does not form a ring itself. (A-B) Sad1-GFP (yellow) and Ppc89-mCherry (magenta) in mitotic kinase mutants (from left to right): wild-type; Aurora B kinase (*ark1-T17*); Cdk1 (*cdc2-asM17*) and Polo kinase (*plo1-24c*). Cells were grown at 25°C or at 36°C for 4 h (WT, *ark1-T17, plo1-24c)* or in the presence of 50 μM DMSO or 1NM-PP1 for 4 h (*cdc2-asM17*) before imaging. (A) Representative SIM Images of early mitotic cells. Inset shows second SPB. (B) Percentage of mitotic cells with a Sad1 ring at the SPB. P values compared to wild- type were determined using the χ^2^ test. All are not significant, except ****, p<0.001. (C) Schematic showing duplicated side-by-side SPBs connected by a bridge on the cytoplasmic face of the ONM. Ppc89 localizes to each SPB core, while Sad1 is present at the INM (Bestul et al. 2017). SPA-SIM of Plo1-GFP (yellow) with Ppc89-mCherry or Sad1-mCherry (magenta) in cells at G2/M using *cdc25.22*. n, number of individual SIM images utilized for averaging. Bars, 500 nm. (D) Representative SIM Images of Plo1- GFP (yellow) and Ppc89-mCherry (magenta) in early mitotic and anaphase cells. Inset shows second SPB in anaphase cells. Bar, 500 nm.

Plo1 has multiple targets that localize throughout the cell, but the kinase is known to localize to the SPB during mitosis (Mulvihill et al., 1999). One particularly tempting idea is that centromere-dependent Polo localization at the SPB initiates Sad1 reorganization and NEBD. This model leads to several testable predictions: Plo1 should localize to the nuclear face of the SPB near Sad1 and the centromere, its localization should be dependent on the SPB-centromere linkage and Polo kinase function should be required for initiation of Sad1 reorganization as well as for all downstream steps, including Cut12 recruitment and NEBD.

High resolution SPA-SIM analysis of Plo1-GFP distribution showed that at the G2/M boundary, the majority of Polo kinase at the SPB is present at the bridge region, which connects the duplicated SPBs marked by Ppc89-mCherry (Fig 5C). The partial co- localization of Plo1-GFP and Sad1-mCherry along with the offset from Ppc89-mCherry indicates that Plo1 is recruited at the INM face of the SPB upon mitotic entry (Fig 5C). This unexpected localization puts the bulk of SPB-localized Plo1 in the vicinity of the centromere and in a location to interact with Sad1. Examination of Plo1-GFP by SPA- SIM in *sad1.2* mutant cells allowed us to examine the role centromere-SPB attachment plays in Polo kinase localization. At 25°C, Plo1-GFP localized to the SPB in almost every cell, giving a sharp averaging signal over background (Fig 5D). Note that this is a distinct pattern of localization compared to wild-type cells (Fig 5C), possibly due to loss/weakened Sad1.2-centromere-attachment even under permissive growth conditions (Fernandez-Alvarez et al., 2016). At 36°C, moderate (4h) or severe (8h) loss of Plo1-GFP at the SPB resulted in diminished Plo1 signal in SPA-SIM (Fig 5D). Given the finding that centromeres become more detached from the SPB after longer periods of incubation at 36°C in *sad1.2* cells (Fernandez-Alvarez et al., 2016), our data showing progressive loss of Plo1-GFP at the SPB in *sad1.2* mutants after incubation at 36°C is consistent with the idea that centromere-SPB attachment is necessary for Plo1 localization to the SPB during prometaphase. However, unlike Sad1, we did not detect ring-like structures of Plo1-GFP at the SPB during any stage of mitosis or in *sad1.2* arrested cells (Fig 5D-E), suggesting that Plo1 does not bind to Sad1 molecules or other components of the mitotic ring in a stoichiometric manner.

To test the requirement for Polo at mitotic onset, it was necessary to generate a synchronized population of cells with and without kinase activity (Fig 6A). Utilizing the *plo1+* analog-sensitive strain, *plo1.as8* (Grallert et al., 2013a), in a *cdc25.22* background, cells were arrested at G2/M by growth at 36°C for 3 h, then the *plo1.as8* inhibitor (3Brb-PP1) was added to inactivate Polo kinase for another 30 min while cells were kept at 36°C. Cultures were then released from the *cdc25.22* arrest by shifting cells to 25°C for 30 min in the presence of 3Brb-PP1 (Fig 6A). In *plo1+* cells or in *plo1.as8* cells treated with vehicle only, mitosis reinitiated within 30 min, but the *plo1.as8* cells treated with 3Brb-PP1 arrest entry due to the requirement for Polo kinase activity. This setup allows us to determine the early mitotic events that require Polo kinase, including its possible role in Sad1 ring formation, Cut12 recruitment and NEBD.

**Figure 6.**
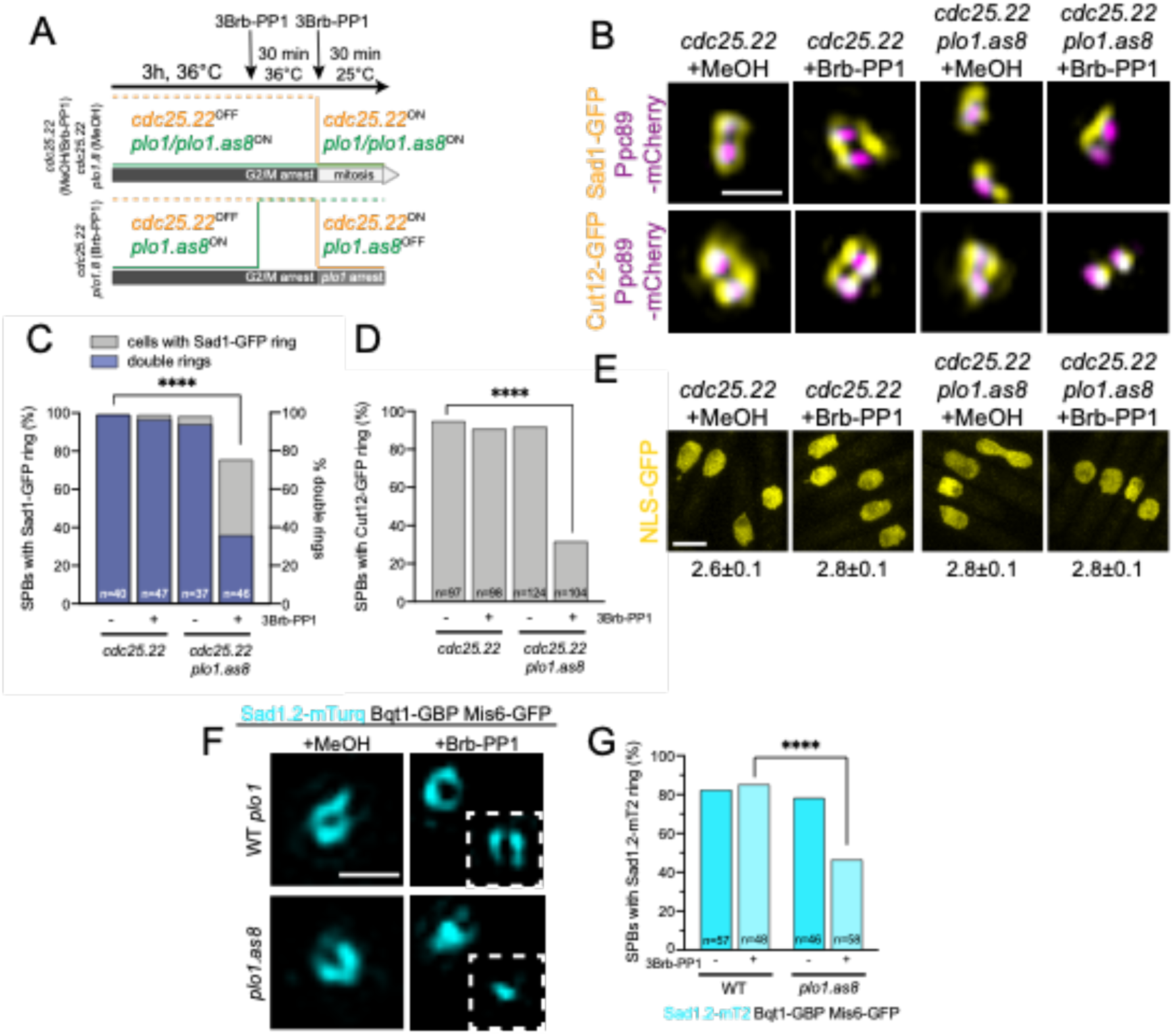
Polo kinase is required for NEBD. (A) To examine if Polo is required at mitotic entry for ring formation, *cdc25.22 plo1.as8* cells were grown for 3 h at 36°C to inactivate *cdc25.22*. Then 3Brb-PP1 was added for 30 min at 36°C to inactivate *plo1.as8*. Shifting cells to 25°C with 3Brb-PP1 for 30 min reactivates *cdc25.22*. In controls, cells enter mitosis, however, inactivation of *plo1.as8* results in a synchronous early mitotic arrest. (B-D) *cdc25.22* or *cdc25.22 plo1.as8* cells with Ppc89-mCherry (magenta) and Sad1-GFP or Cut12-GFP (yellow) were cultured as in Fig 6A. (B) Representative SIM images. Bar, 500 nm. (C) Percentage of mitotic SPBs that had any Sad1-GFP ring (gray) or a double Sad1-GFP ring (blue). (D) Percentage of mitotic SPBs that had any Cut12-GFP ring. P values for C-D compared to wild-type were determined using the χ^2^ test. All are not significant, except ****p<0.001. (E) Nuclear localization of nmt1-3x-NLS-GFP was assayed in *cdc25.22* or *cdc25.22 plo1-as8* to test for NEBD. Cells were grown as in Fig 6A before confocal imaging. Bar, 5 μm. The average ratio of total nuclear to cytoplasmic GFP fluorescence is listed below each image. Error, SEM. n=100. Based on individual t-tests, all differences are not statistically significant. (F-G) Sad1.2-mT2 (cyan) in Bqt1-GBP/Mis6-GFP with wild-type *plo1+* or *plo1.as8* was tested for its ability to form rings by growing cells at 36°C for 8 h in the presence of 50 μM MeOH or 3Brb-PP1 before imaging. (F) Representative SIM image. A second SPB is shown in the inset. Bar, 500 nm. (G) Percentage of cells with Sad1.2-mT2 rings. P values compared to wild-type were determined using the χ^2^ test. All are not significant, except ****p<0.001.

Compared to controls in which virtually all (99%) cells contained a Sad1-GFP ring, 75.6% of cells in which *plo1.as8* was inhibited reorganized Sad1-GFP (Fig 6B-C). Analysis of the rings in control and *plo1.as8* inhibited cells showed a defect in ring maturation in the absence of Polo kinase: only 36% of the rings in *plo1.as8* inhibited cells were mature double rings compared to 95-97% of controls (Fig 6C). This data implies that Polo kinase plays a vital role in Sad1 ring maturation and formation. That we observe a relatively high (75.6%) percentage of *cdc25.22 plo1.8* synchronized cells with Sad1 rings compared to the low (38.5%) percentage seen in *plo1-24c* and other *plo1+* (see Fig 6G; data not shown) mutants suggests that the requirement for Polo in Sad1 ring formation is prior to the *cdc25.22* arrest.

To further define the events downstream of Sad1 ring initiation that might require Polo, we examined Cut12-GFP, which showed a significant reduction in recruitment to rings when Polo kinase activity was inhibited (from 91.9% to 31.7% for Cut12-GFP in *cdc25.22 plo1.as1* cells without and with 3Bbr-PP1) (Fig 6B & D). Thus, Polo acts upstream of Cut12 but downstream of Sad1. Consistent with this idea, we found that *cdc25.22 plo1.as8* cells did not lose NE integrity, as the strain with the inhibitor had the same N:C ratio of GFP fluorescence (2.8±0.1) as without the inhibitor (2.8±0.1) (Fig 6E). Taken together, our data supports a new nuclear role for Polo kinase in the regulation of Sad1 ring maturation and initiation of NEBD during prometaphase.

To confirm that centromere-mediated delivery of Polo kinase to the SPB is needed for Sad1 ring maturation, we triggered Sad1.2-mT2 redistribution with forced centromere binding (Bqt1-GBP, Mis6-GFP) while at the same time inhibiting *plo1.as8* with 3Brb- PP1. Without the analog or in controls, centromere attachment resulted in Sad1.2-mT2 ring formation in ∼80% of cells (Fig 4C-E; 6F-G). However, the fraction of *plo1.as8* strains with Sad1.2-mT2 rings dropped to 46.6% in the presence of the inhibitor (Fig 6G). Examination of individual SPBs showed a full or partial Sad1.2-mT2 ring around one of the two SPBs in the absence of Polo activity, but at the second SPB, ring formation did not occur (Fig 6F). Thus, the centromere itself is insufficient without Polo kinase, which is needed to complete ring formation around both SPBs.

## Discussion

### SPB ring formation at early mitosis facilitates localized NEBD and mitotic progression with Sad1 as the gatekeeper

Utilization of super-resolution microscopy allowed us to visualize a novel mitotic SPB ring surrounding the *S. pombe* SPB for the first time. It contains the NE proteins Sad1 and Kms2, the mitotic regulatory protein Cut12 and the SPB insertion protein Cut11. While a toroidal structure around the SPB has been seen before in *S. cerevisiae* (Chen et al., 2019), the *S. pombe* ring is unique is three ways: 1) it only forms at mitotic entry and dissipates upon mitotic exit; 2) its formation is triggered by the centromere; and 3) Sad1 plays an essential role in ring assembly compared to the important, but non-essential role of Mps3 in *S. cerevisiae* (Chen et al., 2019). The *S. cerevisiae* SPB is inserted into the NE during G1 phase with the help of local NPCs (Rüthnick et al., 2017), which are not observed adjacent to *S. pombe* SPBs (Ding et al., 1997; Uzawa et al., 2004; Tamm et al., 2011). The key role of the LINC complex (Sad1-Kms2/Kms1) in creating the SPB ring could explain how *S. pombe* might coordinate nuclear and cytoplasmic triggers for NE remodeling without NPCs. Importantly, this function of the LINC complex in regulated NEBD could possibly be utilized by other organisms that partially or completely dismantle the NE during mitosis.

Based on our data, we propose the following stepwise model of *S. pombe* ring assembly and NEBD upon mitotic entry (Fig 7). Duplicated SPBs sit on top of the intact NE prior to mitosis surrounded by a large ring of Lem2. During this stage, Sad1 is notpresent in a ring but rather sits at the INM. (1) Upon mitotic entry, Sad1 is redistributed into a ring structure at the INM, mediated in part by centromere interactions. Polo kinase is recruited to the nuclear face of the SPB, although it does not form a ring itself. (2) Further recruitment of Polo kinase leads to redistribution of Kms2 and Cut12, possibly through changes in the LINC complex or the NE. NEBD beneath the SPB is completed. (3) Continued recruitment of Polo kinase via Kms2 to the ONM and via the centromere at the INM results in ring expansion. (4) Lastly, Cut11 is recruited and Lem2 disappears from the SPB ring. This enables the nascent SPBs to insert into the SPB fenestrae and ‘plug the hole’ to prevent complete NEBD.

**Figure 7.**
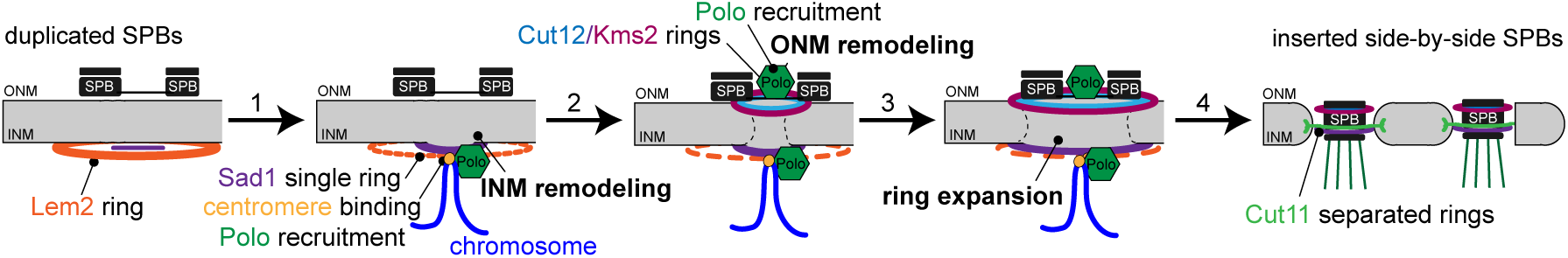
Model of mitotic SPB ring formation and NEBD at *S. pombe* mitotic entry. Duplicated SPBs sit on top of the nuclear envelope with no proteins in a ring structure except Lem2 prior to mitosis. (1) A centromere signal, helped by Csi1, leads to the “old” Sad1 reorganizing into a small ring, opening the INM, while also recruiting Polo. (2) Recruitment of cytoplasmic Polo, Cut12 and Kms2 opens the ONM, completing NEBD. (3) Polo both in the nucleus (modifying the “new” Sad1) and cytoplasm trigger SPB ring expansion to the large ring and encapsulates both SPBs. (4) As Lem2 dissipates, Cut11 is recruited to the mitotic SPBs, allowing SPB insertion and separation.

This timeline provides a snapshot of events leading to NEBD, and it provides a framework from which we can begin to study NEBD at a mechanistic level to address gaps in our model. For example, how is Polo kinase recruited by the centromere? How do Sad1, Kms2 and Cut12 drive NE remodeling? How do Cut11 and Lem2 drive SPB separation? Based on work in other organisms, we can speculate on possible roles for Polo kinase and the LINC complex in NEBD. Following fertilization in *C. elegans*, PLK-1 is recruited by nucleoporins to facilitate localized breakdown of the NE between egg and sperm pronuclei through an unknown mechanism (Martino et al., 2017). In budding yeast, analysis of the Sad1 ortholog Mps3 suggests it plays a role in NEBD by recruitment of proteins involved in lipid remodeling and/or the stabilization of the curved nuclear membrane created at fenestrae (Friederichs et al., 2011; Chen et al., 2019). Yeast two-hybrid analysis suggests that Sad1 may interact with a variety of membrane remodeling factors (Vo et al., 2016; Miki et al., 2004), including Brr6, a protein involved in SPB insertion through its effects on membrane dynamics (Tamm et al., 2011, Zhang et al., 2018).

#### The role of the centromere and centromeric proteins in regulating ring formation

Linkage of chromosomes, specifically telomeres, to the SPB is vital for fission yeast meiosis (Tomita and Cooper, 2007; Klutstein and Cooper, 2014). Although centromeres can substitute for telomeres in meiosis, typically centromere-SPB attachment occurs during mitosis (Fernandez-Alvarez et al., 2016). Here, we show that this connection is needed for SPB ring formation downstream of Sad1 and for NEBD/SPB insertion. Csi1 and Lem2 were tested for their ability to serve as linker proteins between the centromere and Sad1. Although both proteins formed ring-like structures around the SPB, the size and timing of Csi1 rings made it a leading candidate as a Sad1 regulator. Consistent with this idea, Csi1 is known to bind Sad1 and the outer kinetochore protein Spc7 (Hou et al., 2012). Loss of Csi1 function disrupts centromere clustering underneath the SPB (Hou et al., 2012), and Csi1 deletion, when combined with the *sad1.2* mutation causes severe loss of Sad1 ring formation (Fig 4A-B). However, residual Sad1 ring formation in this mutant suggests redundant mechanisms for centromere tethering, possibly through other Sad1-centromere interacting factors or centromere-independent Sad1 regulation.

Lem2 also interacts with Sad1 and the centromere (Hiraoka et al., 2011). Our observation that Lem2 is distributed in a large ring surrounding the SPB during interphase and that its localization to this region dissipates early in mitosis suggests that it only indirectly affects the mitotic SPB, as the *lem2Δ* strain did not affect Sad1 ring formation (Fig S2C). Interestingly, the Lem2 SPB rings completely disappear just before SPB separation at the time when Cut11 localizes to the SPB (Fig S3C). One possibility is that Cut11 replaces Lem2 to drive SPB separation.

Depletion of CENP-A was recently shown to disrupt mitotic spindle formation and displace centrioles (Gemble et al., 2019), a phenotype that is reminiscent of the spindle assembly and disconnected SPBs seen in the *sad1.2* mutant in *S. pombe* (Fernandez-Alvarez et al., 2016). Thus, a key question is how the centromere, centrosome/SPB and spindle coordinate activity to ensure the integrity of the mitotic spindle. We hypothesized that centromeric attachment to the SPB via Sad1 delivered a mitotic regulator to instigate Sad1 reorganization and facilitate NEBD. Using Sad1 ring formation as an assay, we were able to test key mitotic kinases and uncover an unexpected role for Polo kinase.

### Regulation of NEBD through mitotic kinases

In fission yeast, Plo1 is perhaps best known for its role in mitotic activation feedback. Previous work showed that Polo kinase interacts with the SPB components Cut12 and Kms2 to promote mitotic cyclin-Cdk inactivation through the Cdc25 phosphatase and the Wee1 kinase (Walde and King, 2014; Gallert et al., 2013; MacIver et al., 2003). Our data suggest an additional role for Polo kinase, which is likely brought to the nuclear face of the SPB by binding centromere. Our SIM data showing the bulk of Polo at the INM surface of the SPB at the G2/M transition is not inconsistent with data suggesting Plo1 binds to the ONM face of the SPB as our data does not account for the entire population of Plo1 throughout cell division. Interestingly, Sad1 ring formation at one SPB is unaffected by loss of Plo1 while ring formation at the second SPB is blocked (Fig 6F-G). We propose that Plo1 is needed to phosphorylate Sad1 recruited de novo at the ‘new’ SPB, licensing the pole for NE insertion, while the ‘old’ Sad1 was licensed by Plo1 from the previous cell cycle. However, as we have yet to detect direct Plo1 phosphorylation of Sad1 and because Plo1 itself does not form a ring, other intermediates may be involved. Differences in the ‘old’ and ‘new’ SPB have been previously reported in *cut12.1*, *81nmt1-GFP-Kms2*, and *cut11.1* mutants: in each case, the ‘old’ SPB is competent to insert into the NE while the ‘new’ SPB exhibits insertion defects (Bridge et al., 1998; Walde and King 2014; West et al., 1998). In many cases this is accompanied with a loss of NE integrity and loss of a soluble GFP reporter (Tallada et al., 2009; Fernandez-Alvarez et al., 2016; Tamm et al., 2011). Our data brings a new understanding to these phenotypes in several ways. First, the SPB insertion failure is the result of defects in initiation/progression of the mitotic ring cascade. Second, because breakdown of the NE is triggered early in the pathway, mutants defective in SPB insertion will have a loss of NE integrity.

A major question is how Plo1 and the centromere trigger Sad1 reorganization into a ring. Based on EM analysis of SPB structure (Ding et al., 1997; Uzawa et al., 2004), as well as SIM of SPB core components such as Sid4, Pcp1 and Ppc89 (Fig S1A), the redistribution of Sad1 and other ring proteins is not driven by structural changes at the SPB itself. Thus, we presume that Plo1-dependent phosphorylation of Sad1 or another target during prometaphase alters its binding interactions at the SPB to drive ring formation. A leading candidate to facilitate ring formation is Lem2, which is distributed around the SPB throughout interphase. However, as *lem2Δ* cells are still able to form Sad1 rings, other unknown factors are likely involved. Another possibility, which is not mutually exclusive, is that phosphorylation of Sad1 causes it to bind to a protein in a different arrangement that encourages ring formation, similar to Mps3 and Mps2 in budding yeast (Chen et al., 2019). Further analysis of Sad1, its mitotic binding partners and analysis of Plo1 phosphorylation will be required to fully elucidate the mechanism of ring formation.

Somewhat surprisingly, Sad1 ring formation was not dependent on Cdk activity despite having two residues targeted by the kinase (Swaffer MP et al., 2018). The loss of *cdc2+* created more robust Sad1 SPB rings, with significant Sad1 localization away from the SPB (Fig 5A). This suggests mitotic Cdk activity may regulate Sad1 levels in mitotic cells to prevent the formation of ectopic sites of NEBD.

In conclusion, our observations bring to light that redistribution of SPB, NE and centromere proteins in a coordinated fashion is vital to NEBD and mitotic progression. In fission yeast, this process is not regulated by mitotic Cdk but instead it is regulated almost exclusively by the Polo kinase, Plo1. Polo-like kinases facilitate NEBD in *C. elegans* (Rahman et al., 2015) and humans (Lenart et al., 2007), raising the interesting possibility that centromere-linkage, protein redistribution and Polo kinase regulation is more generally involved in nuclear remodeling.

## Materials and Methods

### Yeast Strains and strain construction

*S. pombe* strains used in this study are listed in Table S1, including strains we received from other laboratories: *sad1.2-GFP-KanMX6* and *sad1.2:NatMX6, bleMX6-nmt3X-bqt1- GBP-mCherry:HygMX6, Pnda3-mCherry-atb2:aur1R, Mis6-GFP:kanMX6* (J.P. Cooper, University of Colorado, Denver, CO); *ark1-T7* and *plo1-24c* (K.L. Gould, Vanderbilt University, Nashville, TN); *cdc2-asM17* (K.E. Sawin, University of Edinburgh, Edinburgh, UK); *plo1-GFP-KanMX6* (A. Paoletti, PSL Research University, Paris, France); *plo1.as8-Ura4+* (I.M. Hagan, University of Manchester, Manchester, UK). All fusions to GFP and/or mCherry not listed above were created using PCR-based methods that targeted the endogenous locus as described previously (Bahler et al., 1998). Additional strains were made through standard genetic crosses.

### Cell cycle growth and fixation

To analyze SPB protein distribution at mitotic entry, *cdc25.22* strains with fluorescently tagged SPB components were grown in yeast-extract (YE5S) media for ∼40 h at 25°C, with back dilutions to ensure cells remained logarithmic. Then strains were diluted into Edinburgh minimal media with amino acid supplements (EMM5S) and allowed to grow for 2 h at 25°C before being transferred to 36°C for 3.5 h. Cells were either collected at this time for fixation (see below) or transferred to 25°C for 10, 20 and 30 min before fixation. This growth regimen resulted a homogenous population with good signal-to-noise, which was essential for SIM analysis of single cells. To ensure the *cdc25.22* arrest did not affect protein re-distribution, GFP-tagged SPB proteins were grown in asynchronous wild-type strains and in *cdc25.22* arrested cells shifted to 36°C for 4 h, using previously described growth protocols (P. Fantes 1979) (Fig S1A).

To study *sad1.1*, *cut11.1*, or *cut12.1*, strains were also grown in YE5S media for ∼40 h at 25°C, diluted into EMM5S for 2 h at 25°C, transferred to 36°C for 4 h and then collected for fixation. If strains contained a construct expressed from the *nmt1* promoter (*nmt1-3x-NLS-GFP* and *41nmt1-GFP-Kms2*), growth times and media were altered. Strains were first streaked to EMM5S plates for at least 2 d, then grown in EMM5S media for ∼24 h at 25°C, with back dilutions to ensure cells remained logarithmic. After the 24 h, strains were treated as above for growth and fixation. In separate experiments, these conditions were shown to produce the cell cycle arrest previously reported for each mutant.

To analyze loss of *kms2+* function, *81nmt1-HA-Kms2* strains were streaked out to EMM5S plates for at least 2 d, and then grown in EMM5S media for ∼24 h at 25°C, with back dilutions to ensure cells remained logarithmic. Then 10 µM of thiamine was added to the EMM5S to shut off *kms2+* expression for ∼16 h at 25°C, and then cells were fixed. These conditions recapitulated the cell cycle arrest and loss of HA-Kms2 protein previously reported (Walde and King, 2014).

Analog-sensitive alleles of Polo (*plo1.as8*) and Cdk (*cdc2.asM17*) were inactivated as follows. Cells were grown in YE5S media for ∼40 h at 25°C, with back dilutions to ensure cells remained logarithmic. After diluting into EMM5S for 2 h at 25°C, 50 μM of 1NM-PP1 (Sigma-Aldrich; dissolved in DMSO, *cdc2.asM17*) or 3Brb-PP1 (AbCam; dissolved in methanol, *plo1.as8*) or the vehicle only were added. Cells were then grown for 2 or 4 h before fixation. After 2 h at 25°C in EMM5S, *cdc25.22 plo1.as8* were transferred to 36°C for 3 h before addition of 50 μM 3Brb-PP1/methanol, after which they were allowed to incubated at 36°C for an additional 30 min. Cells were then released from *cdc25.22* by growth at 25°C for 30 min before fixation.

Cells were fixed with 4% paraformaldehyde (Ted Pella) in 100 mM sucrose for 20 min, pelleted by brief centrifugation and then washed twice in PBS, pH 7.4. After the last wash, excess PBS was removed and ∼20 μl of PBS was left to resuspend the cells for visualization by SIM. While fixation was not required to visualize any protein distribution reported here, it aided SIM analysis by immobilizing the SPB and ensuring protein distribution was not affected by room temperature imaging.

### SIM imaging and SPA-SIM

SIM imaging utilized an Applied Precision OMX Blaze V4 (GE Healthcare) using a 60x 1.42 NA Olympus Plan Apo oil objective and two PCO Edge sCMOS cameras (one camera for each channel). All SIM microscopy was performed at room temperature (22°C-23°C). For the two-color GFP/mCherry experiments, a 405/488/561/640 dichroic was used with 504- to 552-nm and 590- to 628-nm emission filters for GFP and mCherry, respectively. Images were taken using a 488-nm laser (for GFP) or a 561-nm laser (for mCherry), with alternating excitation. SIM reconstruction was done with Softworx (Applied Precision Ltd.) with a Wiener filter of 0.001. SIM images shown in the publication are maximum projections of all z-slices, scaled 8 x 8 with bilinear interpolation using ImageJ (National Institutes of Health) to enlarge the images.

SPA-SIM analysis was performed with custom written macros and plugins in ImageJ. All plugins and source code are available at http://research.stowers.org/imagejplugins/. Individual spots of mother and satellite SPBs were fitted to two 3D Gaussian functions and realigned along the axis between these functions for further analysis using [jay_sim_fitting_macro_multicolor_profile_NPC.ijm]. Spot selection was performed in a semiautomated fashion with manual identification and selection of mother and daughter SPBs. A secondary protein (Ppc89-mCherry) was used as a fiduciary marker to determine position of the GFP-labeled protein so that all positions of the SPB proteins were compared with a single origin point. For the fiducial protein, the higher intensity spot was assigned as the mother SPB. After alignment, images were averaged and scaled as described previously (Burns et al., 2015), using [merge_all_stacks_jru_v1.ijm] then [stack_statistics_jru_v2.ijm].

To quantitate the distribution of GFP-tagged SPB proteins and assess ring formation, images containing the GFP-tagged protein and Ppc89-mCherry were used. Individual images were manually inspected and analyzed as follows. If the GFP signal encompassed over 50% of the Ppc89-mCherry signal, it was counted as a ring. In some cases, the ring was in the z-axis (thus not visible as a ring in the xy-plane); if the GFP signal extended beyond the Ppc89-mCherry signal for more than 50 nm in both directions, then it was also tabulated as a ring. Because of the small size of some rings and the limited resolution of SIM (particularly in the z-axis), not all rings had a distinct center.

### Confocal imaging and analysis

Confocal imaging utilized a PerkinElmer UltraVIEW VoX with a Yokogawa CSU-X1 spinning disk head, a 100x 1.46 NA Olympus Plan Apo oil objective and CCD (ORCA- R2) and EMCCD (C9100-13) cameras. All confocal microscopy was performed at room temperature (22°C-23°C). GFP/mCherry images were taken using a 488-nm laser (for GFP) or a 561-nm laser (for mCherry), with alternating excitation. Images were collected using the Volocity imaging software.

To measure the intensity of NLS-GFP, sum projections of the entire z-stack were created using ImageJ. After background subtraction, a region of interest (ROI) was drawn around each nucleus and the integrated fluorescence intensity was divided by the area of the ROI. This ROI was then used to measure the integrated fluorescence intensity of the cytoplasm of that cell. The integrated fluorescence intensity of the nucleus over the cytoplasm gave a N:C ratio for each cell. The average of this ratio for 100 cells was determined.

## Acknowledgements

We are grateful to Julie Cooper, Kathy Gould, Ken Sawin, Anne Paoletti and Iain Hagan for reagents and to the Jaspersen Lab for discussions and comments on the manuscript. Research reported in this publication was supported by the Stowers Institute for Medical Research and the NIH-NIGMS under award number R01GM121443 (to SLJ). Original data underlying this manuscript can be downloaded from the Stowers Original Data Repository at http://www.stowers.org/research/publications/LIBPB-xxxx. The authors declare no competing financial interests.

## Author Contributions

AJB and SLJ conceived the experiments, AJB constructed strains and performed experiments, ZU assisted with SIM, JRU developed tools for image analysis, AJB prepared figures, and AJB and SLJ wrote the paper with input from all the authors.

## Abbreviations

MTOC: microtubule-organizing center
SPB: spindle pole body
SIM: structured illumination microscopy
EM: electron microscopy
NE: nuclear envelope
NPC: nuclear pore complex
SPA: single particle analysis
SEM: standard error of the mean
EM: electron microscopy
SPIN: spindle pole body insertion network
SUN: Sad1-UNC-84 homology
KASH: Klarsicht-ANC-1-Syne-1 homology
LINC: linker of nucleoskeleton and cytoskeleton
mT2: mTurquoise2
NEBD: nuclear envelope break down
CDK: cyclin-dependent kinase
ROI: region of interest

## Supplementary Material

**Table S1.**
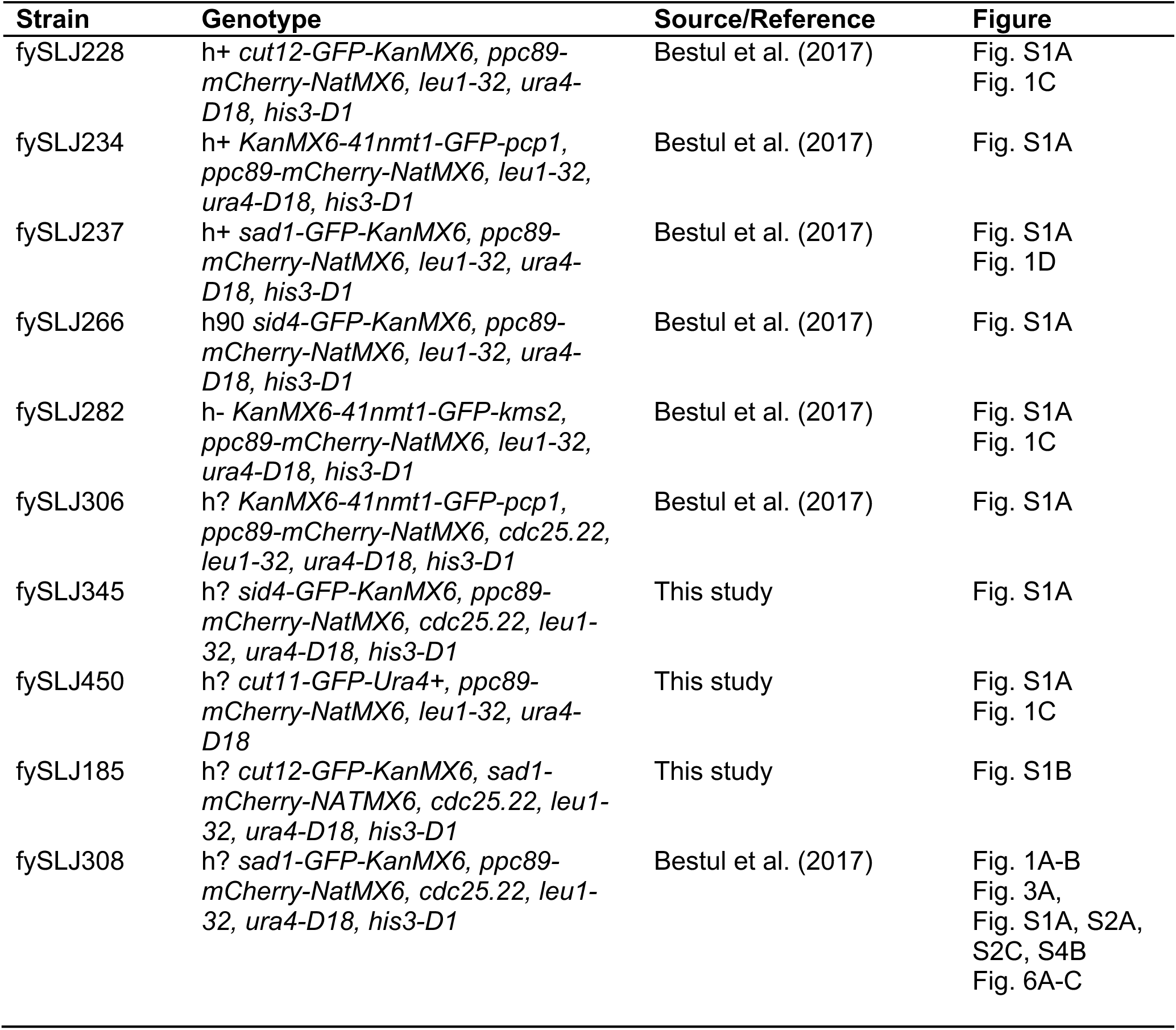
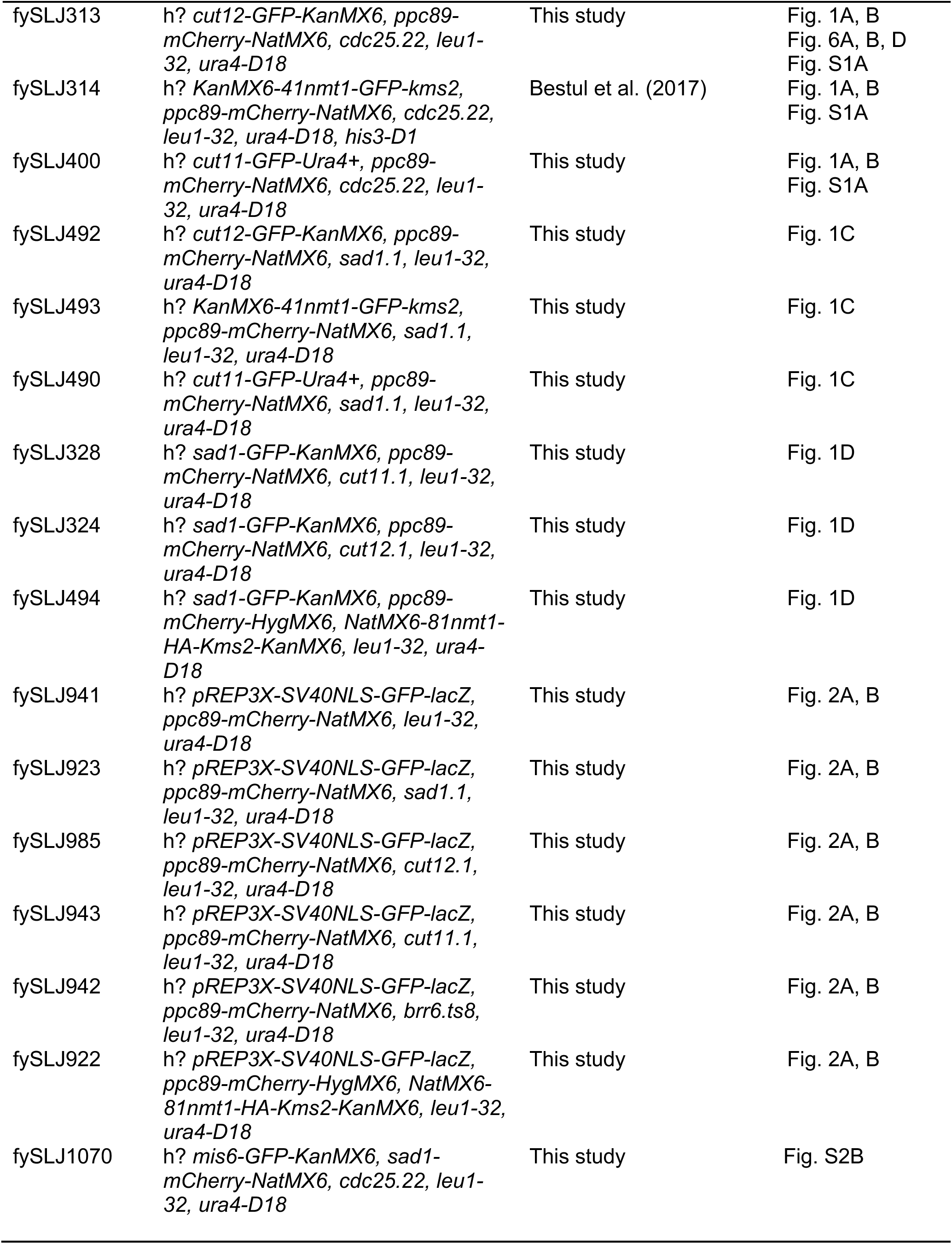
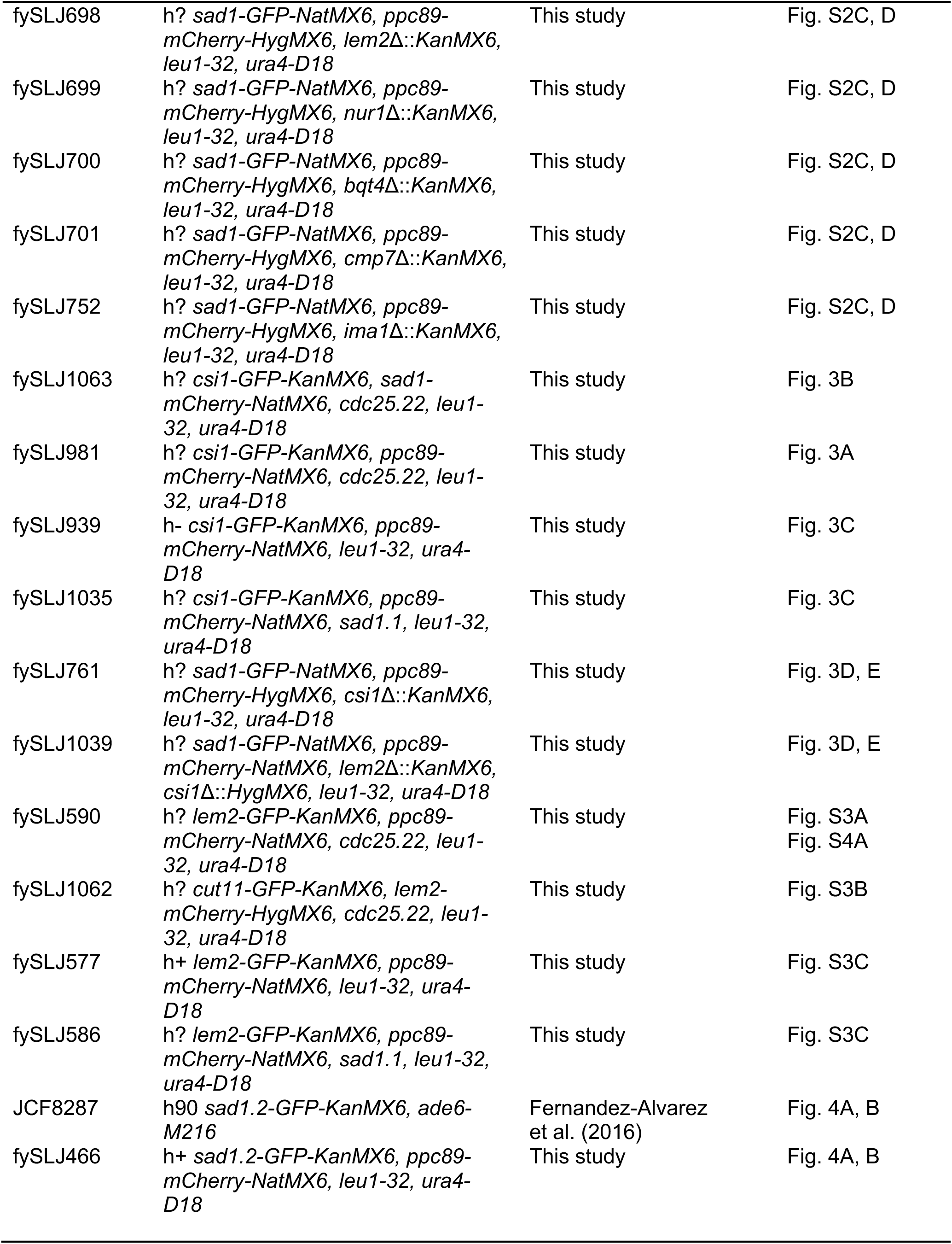
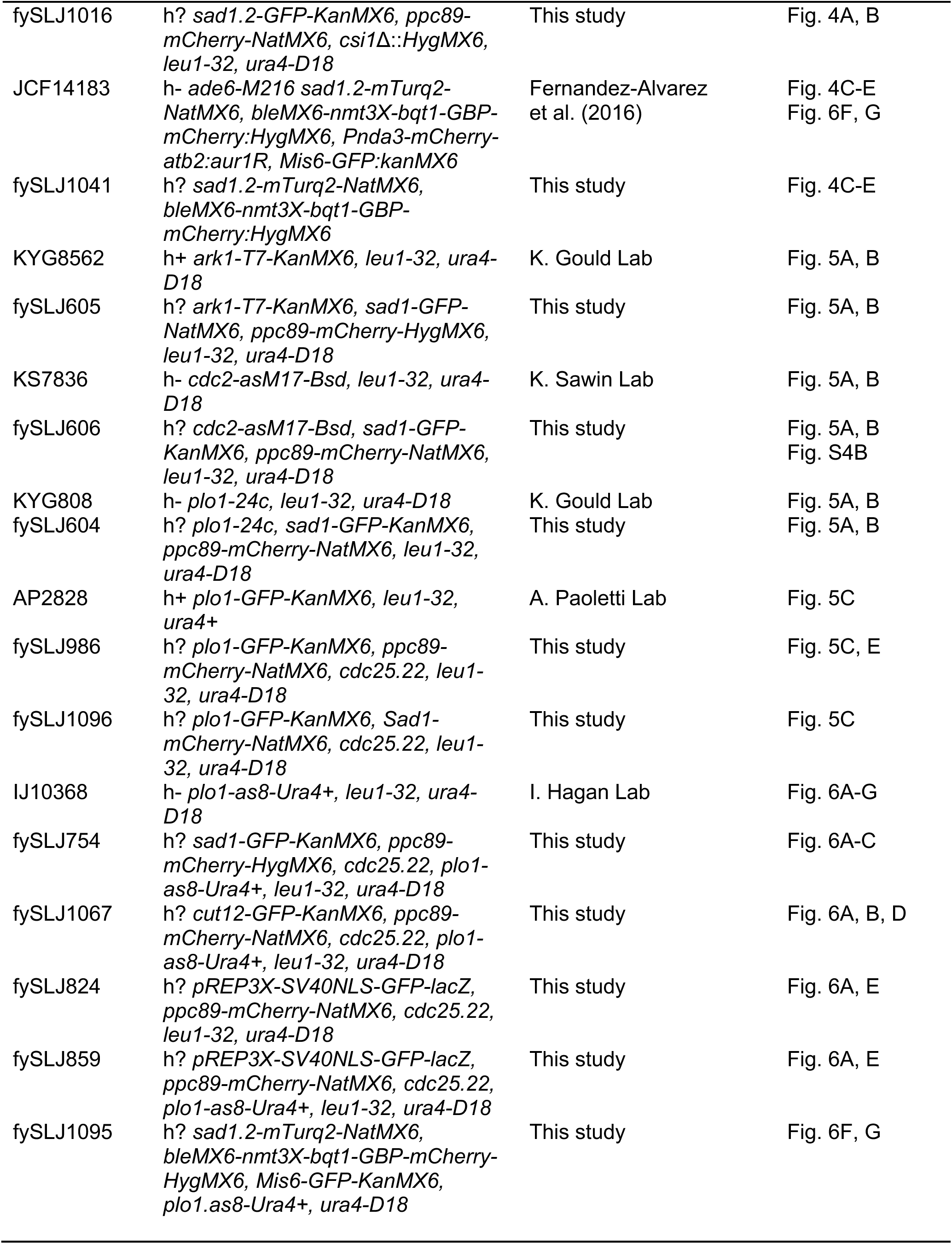
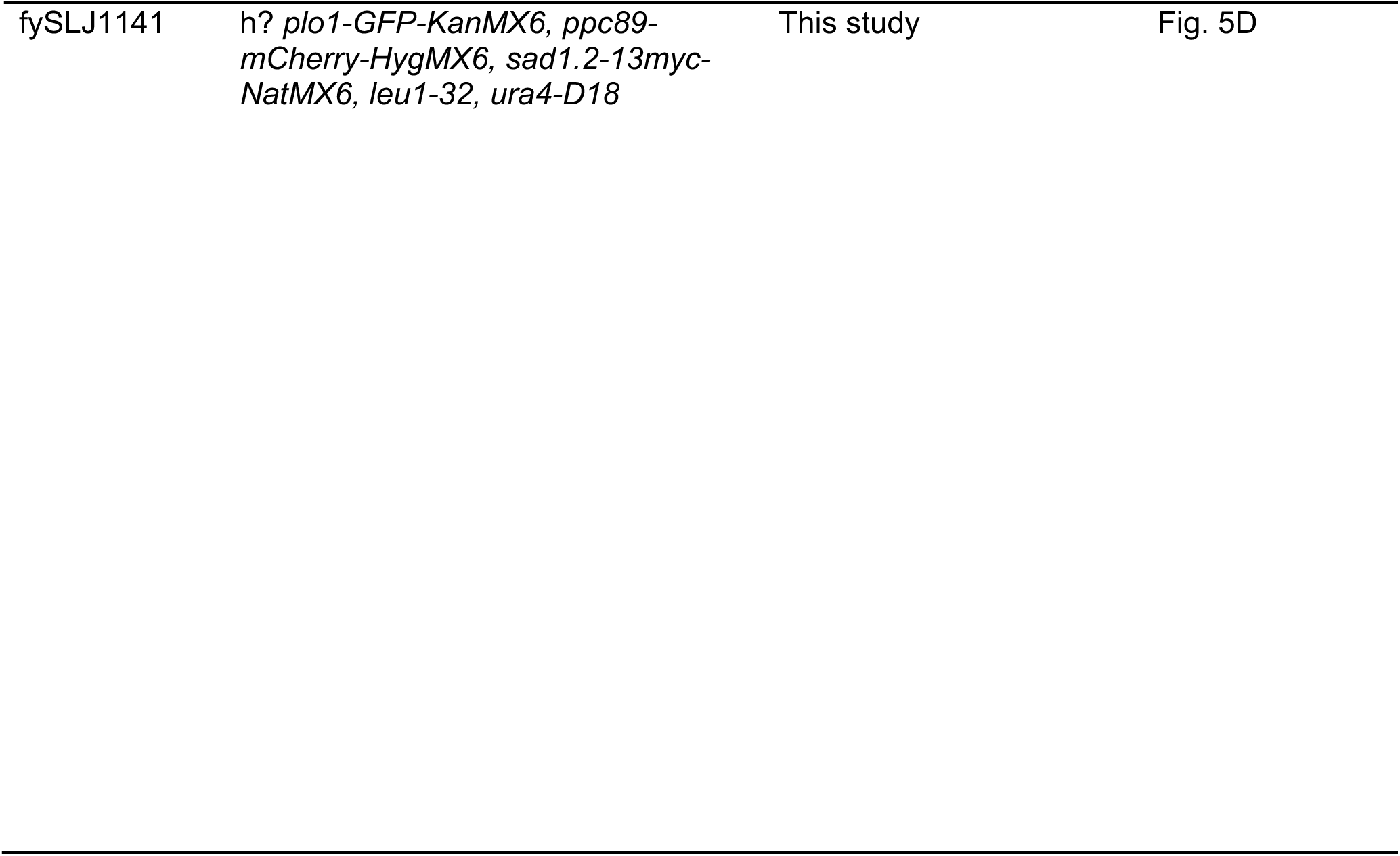
Yeast Strains.

| **Strain** | **Genotype** | **Source/Reference** | **Figure** |
| --- | --- | --- | --- |
| fySLJ228 | h+ *cut12-GFP-KanMX6, ppc89-mCherry-NatMX6, leu1-32, ura4-D18, his3-D1* | Bestul et al. (2017) | Fig. S1A<br>Fig. 1C |
| fySLJ234 | h+ *KanMX6-41nmt1-GFP-pcp1, ppc89-mCherry-NatMX6, leu1-32, ura4-D18, his3-D1* | Bestul et al. (2017) | Fig. S1A |
| fySLJ237 | h+ *sad1-GFP-KanMX6, ppc89-mCherry-NatMX6, leu1-32, ura4-D18, his3-D1* | Bestul et al. (2017) | Fig. S1A<br>Fig. 1D |
| fySLJ266 | h90 *sid4-GFP-KanMX6, ppc89-mCherry-NatMX6, leu1-32, ura4-D18, his3-D1* | Bestul et al. (2017) | Fig. S1A |
| fySLJ282 | h- *KanMX6-41nmt1-GFP-kms2, ppc89-mCherry-NatMX6, leu1-32, ura4-D18, his3-D1* | Bestul et al. (2017) | Fig. S1A<br>Fig. 1C |
| fySLJ306 | h? *KanMX6-41nmt1-GFP-pcp1, ppc89-mCherry-NatMX6, cdc25.22, leu1-32, ura4-D18, his3-D1* | Bestul et al. (2017) | Fig. S1A |
| fySLJ345 | h? *sid4-GFP-KanMX6, ppc89-mCherry-NatMX6, cdc25.22, leu1-32, ura4-D18, his3-D1* | This study | Fig. S1A |
| fySLJ450 | h? *cut11-GFP-Ura4+, ppc89-mCherry-NatMX6, leu1-32, ura4-D18* | This study | Fig. S1A<br>Fig. 1C |
| fySLJ185 | h? *cut12-GFP-KanMX6, sad1-mCherry-NATMX6, cdc25.22, leu1-32, ura4-D18, his3-D1* | This study | Fig. S1B |
| fySLJ308 | h? *sad1-GFP-KanMX6, ppc89-mCherry-NatMX6, cdc25.22, leu1-32, ura4-D18, his3-D1* | Bestul et al. (2017) | Fig. 1A-B<br>Fig. 3A,<br>Fig. S1A, S2A,<br>S2C, S4B<br>Fig. 6A-C |
| fySLJ313 | h? *cut12-GFP-KanMX6, ppc89-mCherry-NatMX6, cdc25.22, leu1-32, ura4-D18* | This study | Fig. 1A, B<br>Fig. 6A, B, D<br>Fig. S1A |
| fySLJ314 | h? *KanMX6-41nmt1-GFP-kms2, ppc89-mCherry-NatMX6, cdc25.22, leu1-32, ura4-D18, his3-D1* | Bestul et al. (2017) | Fig. 1A, B<br>Fig. S1A |
| fySLJ400 | h? *cut11-GFP-Ura4+, ppc89-mCherry-NatMX6, cdc25.22, leu1-32, ura4-D18* | This study | Fig. 1A, B<br>Fig. S1A |
| fySLJ492 | h? *cut12-GFP-KanMX6, ppc89-mCherry-NatMX6, sad1.1, leu1-32, ura4-D18* | This study | Fig. 1C |
| fySLJ493 | h? *KanMX6-41nmt1-GFP-kms2, ppc89-mCherry-NatMX6, sad1.1, leu1-32, ura4-D18* | This study | Fig. 1C |
| fySLJ490 | h? *cut11-GFP-Ura4+, ppc89-mCherry-NatMX6, sad1.1, leu1-32, ura4-D18* | This study | Fig. 1C |
| fySLJ328 | h? *sad1-GFP-KanMX6, ppc89-mCherry-NatMX6, cut11.1, leu1-32, ura4-D18* | This study | Fig. 1D |
| fySLJ324 | h? *sad1-GFP-KanMX6, ppc89-mCherry-NatMX6, cut12.1, leu1-32, ura4-D18* | This study | Fig. 1D |
| fySLJ494 | h? *sad1-GFP-KanMX6, ppc89-mCherry-HygMX6, NatMX6-81nmt1-HA-Kms2-KanMX6, leu1-32, ura4-D18* | This study | Fig. 1D |
| fySLJ941 | h? *pREP3X-SV40NLS-GFP-lacZ, ppc89-mCherry-NatMX6, leu1-32, ura4-D18* | This study | Fig. 2A, B |
| fySLJ923 | h? *pREP3X-SV40NLS-GFP-lacZ, ppc89-mCherry-NatMX6, sad1.1, leu1-32, ura4-D18* | This study | Fig. 2A, B |
| fySLJ985 | h? *pREP3X-SV40NLS-GFP-lacZ, ppc89-mCherry-NatMX6, cut12.1, leu1-32, ura4-D18* | This study | Fig. 2A, B |
| fySLJ943 | h? *pREP3X-SV40NLS-GFP-lacZ, ppc89-mCherry-NatMX6, cut11.1, leu1-32, ura4-D18* | This study | Fig. 2A, B |
| fySLJ942 | h? *pREP3X-SV40NLS-GFP-lacZ, ppc89-mCherry-NatMX6, brr6.ts8, leu1-32, ura4-D18* | This study | Fig. 2A, B |
| fySLJ922 | h? *pREP3X-SV40NLS-GFP-lacZ, ppc89-mCherry-HygMX6, NatMX6-81nmt1-HA-Kms2-KanMX6, leu1-32, ura4-D18* | This study | Fig. 2A, B |
| fySLJ1070 | h? *mis6-GFP-KanMX6, sad1-mCherry-NatMX6, cdc25.22, leu1-32, ura4-D18* | This study | Fig. S2B |
| fySLJ698 | h? *sad1*-GFP-NatMX6, *ppc89*-*mCherry*-HygMX6, *lem2Δ::KanMX6*, *leu1-32*, *ura4-D18* | This study | Fig. S2C, D |
| fySLJ699 | h? *sad1*-GFP-NatMX6, *ppc89*-*mCherry*-HygMX6, *nur1Δ::KanMX6*, *leu1-32*, *ura4-D18* | This study | Fig. S2C, D |
| fySLJ700 | h? *sad1*-GFP-NatMX6, *ppc89*-*mCherry*-HygMX6, *bqt4Δ::KanMX6*, *leu1-32*, *ura4-D18* | This study | Fig. S2C, D |
| fySLJ701 | h? *sad1*-GFP-NatMX6, *ppc89*-*mCherry*-HygMX6, *cmp7Δ::KanMX6*, *leu1-32*, *ura4-D18* | This study | Fig. S2C, D |
| fySLJ752 | h? *sad1*-GFP-NatMX6, *ppc89*-*mCherry*-HygMX6, *ima1Δ::KanMX6*, *leu1-32*, *ura4-D18* | This study | Fig. S2C, D |
| fySLJ1063 | h? *csi1*-GFP-KanMX6, *sad1*-*mCherry*-NatMX6, *cdc25.22*, *leu1-32*, *ura4-D18* | This study | Fig. 3B |
| fySLJ981 | h? *csi1*-GFP-KanMX6, *ppc89*-*mCherry*-NatMX6, *cdc25.22*, *leu1-32*, *ura4-D18* | This study | Fig. 3A |
| fySLJ939 | h- *csi1*-GFP-KanMX6, *ppc89*-*mCherry*-NatMX6, *leu1-32*, *ura4-D18* | This study | Fig. 3C |
| fySLJ1035 | h? *csi1*-GFP-KanMX6, *ppc89*-*mCherry*-NatMX6, *sad1.1*, *leu1-32*, *ura4-D18* | This study | Fig. 3C |
| fySLJ761 | h? *sad1*-GFP-NatMX6, *ppc89*-*mCherry*-HygMX6, *csi1Δ::KanMX6*, *leu1-32*, *ura4-D18* | This study | Fig. 3D, E |
| fySLJ1039 | h? *sad1*-GFP-NatMX6, *ppc89*-*mCherry*-NatMX6, *lem2Δ::KanMX6*, *csi1Δ::HygMX6*, *leu1-32*, *ura4-D18* | This study | Fig. 3D, E |
| fySLJ590 | h? *lem2*-GFP-KanMX6, *ppc89*-*mCherry*-NatMX6, *cdc25.22*, *leu1-32*, *ura4-D18* | This study | Fig. S3A<br>Fig. S4A |
| fySLJ1062 | h? *cut11*-GFP-KanMX6, *lem2*-*mCherry*-HygMX6, *cdc25.22*, *leu1-32*, *ura4-D18* | This study | Fig. S3B |
| fySLJ577 | h+ *lem2*-GFP-KanMX6, *ppc89*-*mCherry*-NatMX6, *leu1-32*, *ura4-D18* | This study | Fig. S3C |
| fySLJ586 | h? *lem2*-GFP-KanMX6, *ppc89*-*mCherry*-NatMX6, *sad1.1*, *leu1-32*, *ura4-D18* | This study | Fig. S3C |
| JCF8287 | h90 *sad1.2*-GFP-KanMX6, *ade6-M216* | Fernandez-Alvarez et al. (2016) | Fig. 4A, B |
| fySLJ466 | h+ *sad1.2*-GFP-KanMX6, *ppc89*-*mCherry*-NatMX6, *leu1-32*, *ura4-D18* | This study | Fig. 4A, B |
| fySLJ1016 | h? *sad1.2-GFP-KanMX6, ppc89-mCherry-NatMX6, csi1Δ::HygMX6, leu1-32, ura4-D18* | This study | Fig. 4A, B |
| JCF14183 | h- *ade6-M216 sad1.2-mTurq2-NatMX6, bleMX6-nmt3X-bqt1-GBP-mCherry:HygMX6, Pnda3-mCherry-atb2:aur1R, Mis6-GFP:kanMX6* | Fernandez-Alvarez et al. (2016) | Fig. 4C-E<br>Fig. 6F, G |
| fySLJ1041 | h? *sad1.2-mTurq2-NatMX6, bleMX6-nmt3X-bqt1-GBP-mCherry:HygMX6* | This study | Fig. 4C-E |
| KYG8562 | h+ *ark1-T7-KanMX6, leu1-32, ura4-D18* | K. Gould Lab | Fig. 5A, B |
| fySLJ605 | h? *ark1-T7-KanMX6, sad1-GFP-NatMX6, ppc89-mCherry-HygMX6, leu1-32, ura4-D18* | This study | Fig. 5A, B |
| KS7836 | h- *cdc2-asM17-Bsd, leu1-32, ura4-D18* | K. Sawin Lab | Fig. 5A, B |
| fySLJ606 | h? *cdc2-asM17-Bsd, sad1-GFP-KanMX6, ppc89-mCherry-NatMX6, leu1-32, ura4-D18* | This study | Fig. 5A, B<br>Fig. S4B |
| KYG808 | h- *plo1-24c, leu1-32, ura4-D18* | K. Gould Lab | Fig. 5A, B |
| fySLJ604 | h? *plo1-24c, sad1-GFP-KanMX6, ppc89-mCherry-NatMX6, leu1-32, ura4-D18* | This study | Fig. 5A, B |
| AP2828 | h+ *plo1-GFP-KanMX6, leu1-32, ura4+* | A. Paoletti Lab | Fig. 5C |
| fySLJ986 | h? *plo1-GFP-KanMX6, ppc89-mCherry-NatMX6, cdc25.22, leu1-32, ura4-D18* | This study | Fig. 5C, E |
| fySLJ1096 | h? *plo1-GFP-KanMX6, Sad1-mCherry-NatMX6, cdc25.22, leu1-32, ura4-D18* | This study | Fig. 5C |
| IJ10368 | h- *plo1-as8-Ura4+, leu1-32, ura4-D18* | I. Hagan Lab | Fig. 6A-G |
| fySLJ754 | h? *sad1-GFP-KanMX6, ppc89-mCherry-HygMX6, cdc25.22, plo1-as8-Ura4+, leu1-32, ura4-D18* | This study | Fig. 6A-C |
| fySLJ1067 | h? *cut12-GFP-KanMX6, ppc89-mCherry-NatMX6, cdc25.22, plo1-as8-Ura4+, leu1-32, ura4-D18* | This study | Fig. 6A, B, D |
| fySLJ824 | h? *pREP3X-SV40NLS-GFP-lacZ, ppc89-mCherry-NatMX6, cdc25.22, leu1-32, ura4-D18* | This study | Fig. 6A, E |
| fySLJ859 | h? *pREP3X-SV40NLS-GFP-lacZ, ppc89-mCherry-NatMX6, cdc25.22, plo1-as8-Ura4+, leu1-32, ura4-D18* | This study | Fig. 6A, E |
| fySLJ1095 | h? *sad1.2-mTurq2-NatMX6, bleMX6-nmt3X-bqt1-GBP-mCherry-HygMX6, Mis6-GFP-KanMX6, plo1.as8-Ura4+, ura4-D18* | This study | Fig. 6F, G |
| fySLJ1141 | h? *plo1-GFP-KanMX6*, *ppc89-mCherry-HygMX6*, *sad1.2-13myc-NatMX6*, *leu1-32*, *ura4-D18* | This study | Fig. 5D |

**Figure S1.**
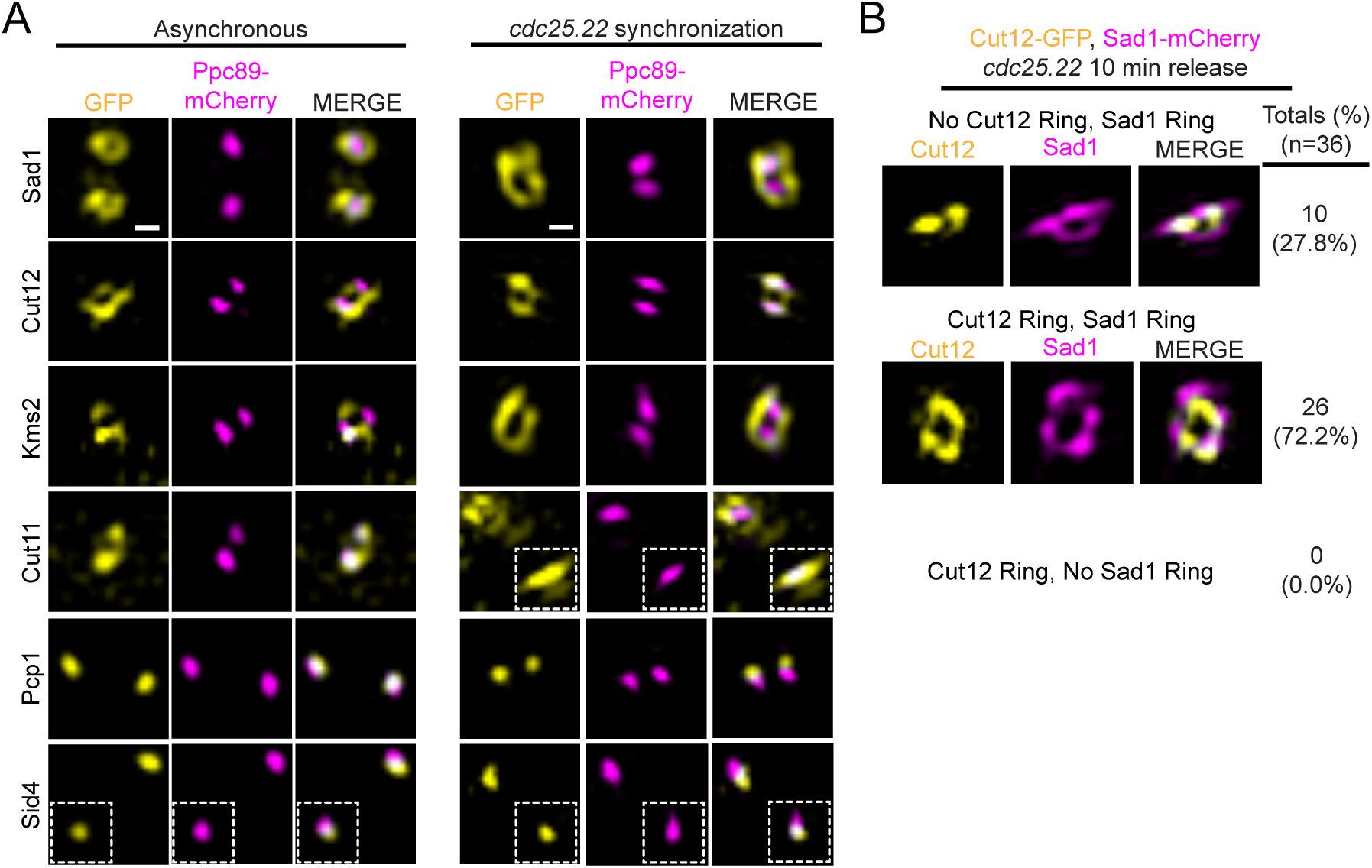
Four specific SPB proteins form a mitotic ring independent of cell cycle synchronization. (A) Asynchronous wild-type cells (left column) or *cdc25.22* cells (right column) containing Ppc89-mCherry (magenta) and the indicated SPB protein fused to GFP (yellow). Wild-type cells were grown for 4 h at 25°C before imaging by SIM. *cdc25.22* cells were grown for 4 h at 36°C and then released to 25°C for 15 min before imaging by SIM. Insets show the second SPB present in the same cell. (B) *cdc25.22* cells containing Sad1-mCherry (magenta) and Cut12-GFP (yellow) were grown for 3.5 h at 36°C and then released to 25°C for 10 min before imaging by SIM. Mitotic ring structures were binned into 3 categories, as illustrated in the example images: 1) No Cut12 ring, Sad1 ring; 2) Both Cut12 and Sad1 ring; 3) Cut12 ring; No Sad1 ring (not observed). The percentage of each is indicated, along with the n-values.

**Figure S2.**
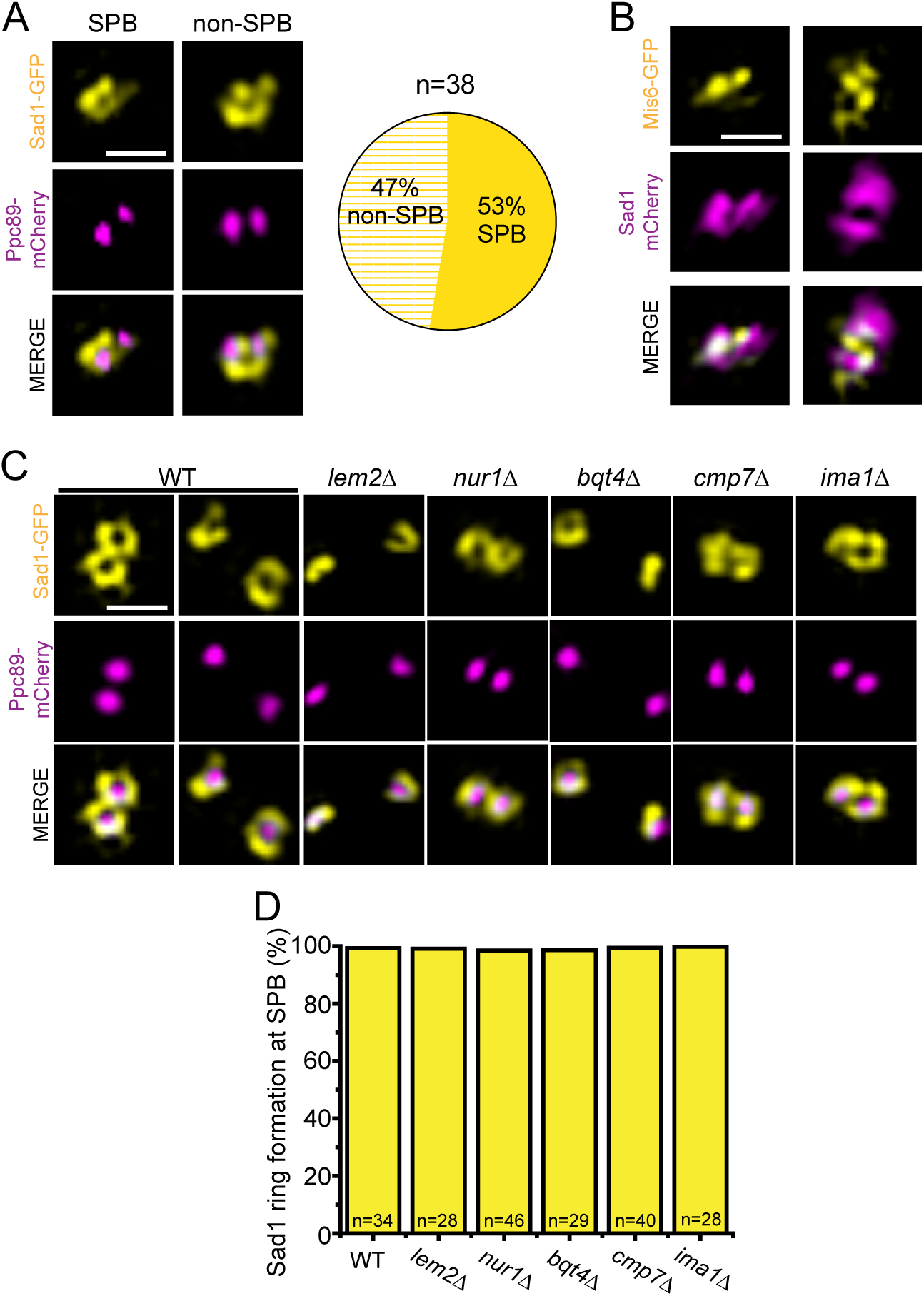
Sad1 ring formation occurs in proximity to the centromeres. (A) *cdc25.22* cells containing Ppc89-mCherry (magenta) and Sad1-GFP (yellow) were grown for 3.5 h at 36°C and then released to 25°C for 10 min before imaging by SIM. The Sad1-GFP ring co-localized with the SPB (left) or to an area in between the two SPBs (right, non-SPB). The percentage of cells in each configuration is shown. (B) Similarly, *cdc25.22* cells with Sad1-mCherry (magenta) and Mis6-GFP (yellow) were arrested and released to determine the position of the centromere relative to the Sad1 ring. (C-D) *cdc25.22* cells containing Ppc89-mCherry (magenta) and Sad1-GFP (yellow) in the indicated centromere/INM protein deletion background were arrested at the G2/M boundary and analyzed by SIM. (C) Representative SIM images. (D) Percentage of mitotic SPBs with Sad1-GFP rings. Bars, 500 nm.

**Figure S3.**
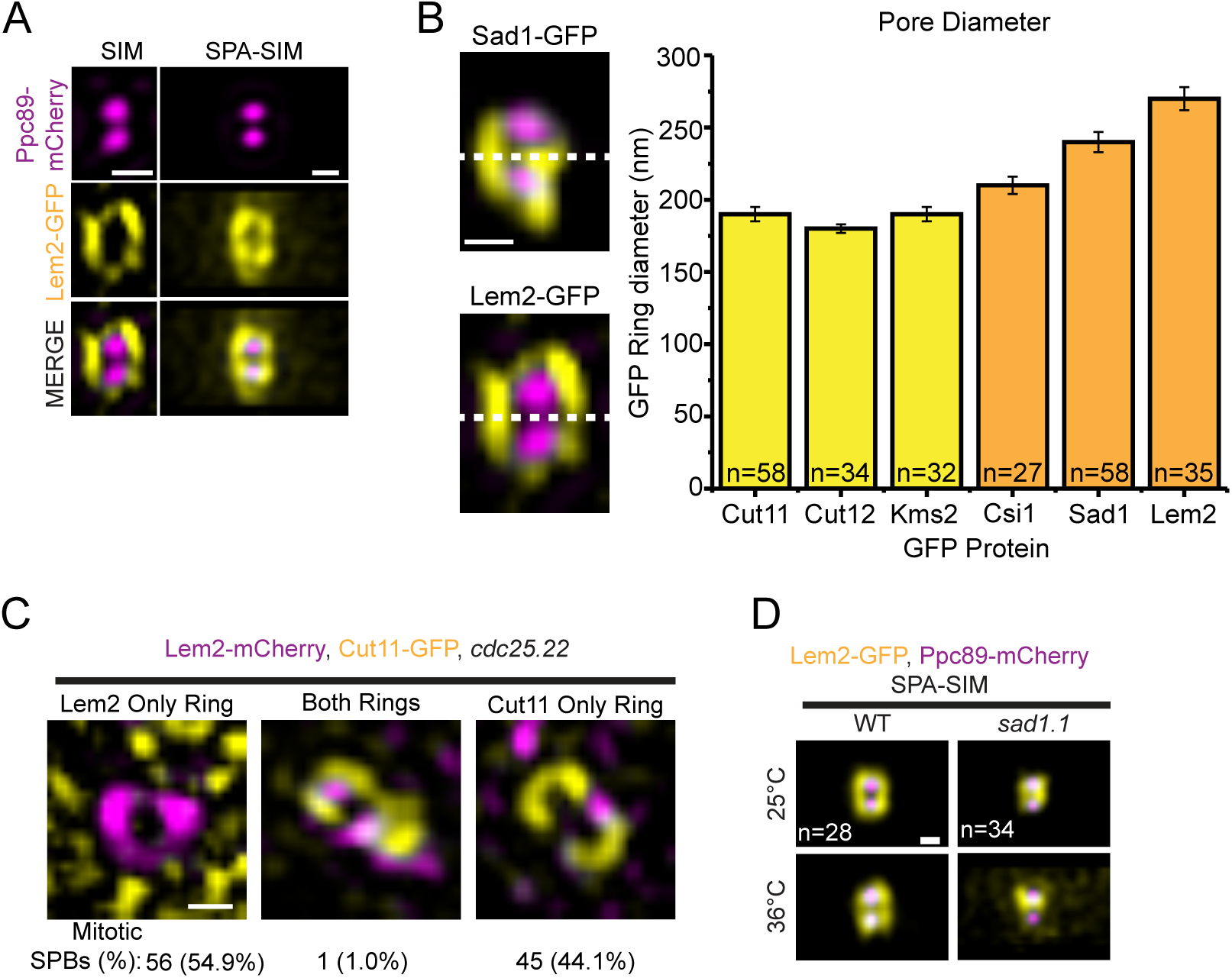
Lem2 forms a unique ring during interphase and early mitosis. (A) *cdc25.22* arrested cells containing Lem2-GFP (yellow) and Ppc89-mCherry (magenta) were imaged by SIM. Individual SIM image (left) and SPA-SIM (right) from the indicated number of images. Bar, 200 nm. (B) To determine the size of SPB rings, a five-pixel wide line was drawn across the ring in between the two SPBs in individual SIM images from *cdc25.22* arrested cells containing the indicated GFP-tagged protein (yellow) and Ppc89-mCherry (magenta), as shown for the examples of Sad1-GFP and Lem2-GFP. Bar, 200 nm. The difference in peak intensity along the line was used as the ring diameter (in nm). Average ring diameter for each protein is plotted, with yellow bars showing similar, smaller diameters and orange bars indicating proteins with larger diameters. Error bars, SEM. (C) *cdc25.22* arrested cells containing Lem2-mCherry (magenta) and Cut11-GFP (yellow) were released into mitosis at 25°C for 30 min and then imaged by SIM. Three configurations were observed in the indicated fraction of cells: Lem2 ring with no Cut11 ring (left), Cut11 ring with no Lem2 ring (center), and rings of both Cut11 and Lem2 (right). n=102. Bar, 200 nm. (C) SPA-SIM images for Lem2-GFP in wild-type and *sad1.1* cells grown at 25°C or 36°C for 4 h. n, number of individual SIM images utilized for averaging. Bar, 200 nm.

**Figure S4.**
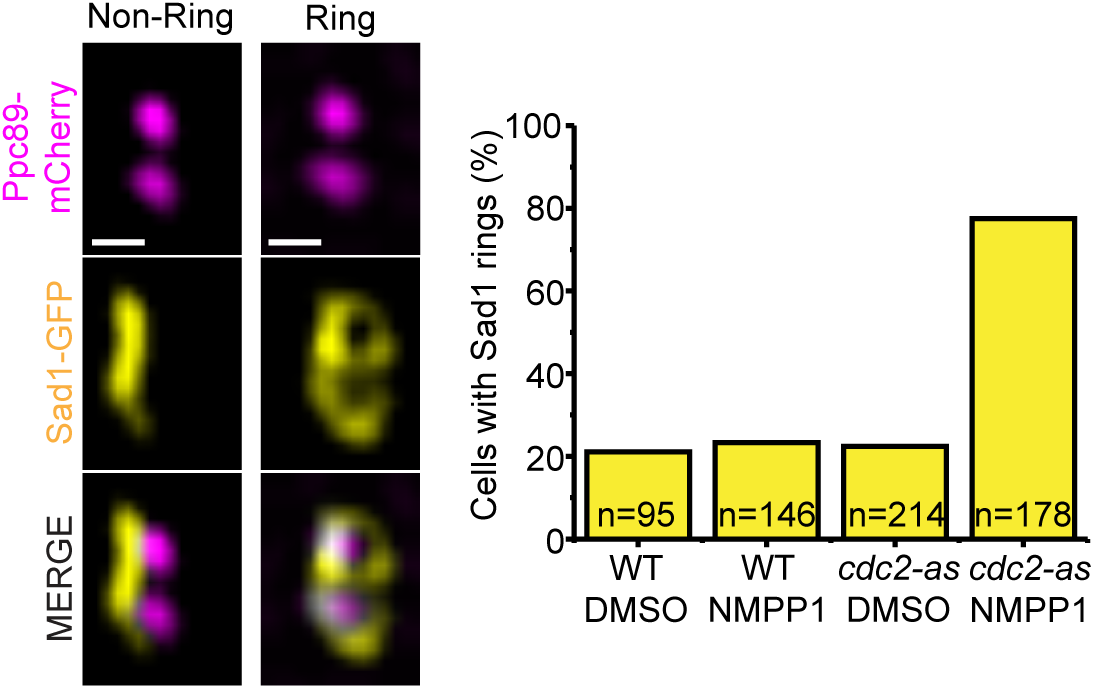
Cdc2 inhibition results in an early mitotic arrest after Sad1 ring formation. Wild-type and *cdc2.asM17* cells containing Sad1-GFP (yellow) and Ppc89- mCherry (magenta) were grown at 25°C in the presence of 50 μM DMSO or 1NM-PP1 for 4 h then imaged by SIM. These cells were scored for the presence (ring) or absence (non-ring) of a Sad1-GFP ring around the SPB(s), as shown in the example. Bar, 200 nm. Typically, in cycling cells, ∼20% of SPBs are in a mitotic configuration with Sad1- GFP visible in rings. When *cdc2.asM17* is inhibited with 1NM-PP1, the fraction of mitotic cells is 77.5%. Because SPBs are not separated, we infer that these cells are stalled in prometaphase before SPB separation.

